# 3’ UTR Insertion of a Directed-Evolved RNA Element for Enhanced Translation

**DOI:** 10.64898/2026.05.07.723449

**Authors:** Xiyang Liu, Qinhao Zhang, Jing Wang, Zheping Zhang, Liqin Zhang

**Author notes:** These authors contributed equally (Xiyang Liu, Qinhao Zhang).

## Abstract

Translation efficiency remains a major limitation for RNA therapeutics. Conventional optimization targets the 5’ untranslated region (5’ UTR), while the 3’ UTR is viewed mainly as a stabilizing element. Here, we demonstrate that the 3’ UTR can be rationally engineered to actively enhance translation. Using an intracellular directed-evolution platform based on the SINEB2 element, we identified RNA modules P51 and its compact variant P51t3,which markedly increased protein output without affecting mRNA levels. P51t3 consistently boosted expression two- to six-fold across plasmid, in vitro transcribed mRNA, and recombinant AAV systems. Mechanistic studies revealed that P51t3 binds ribosomal protein RPL39, recruiting 60S subunits to the initiation site through the natural closed-loop translation model. By integrating evolutionary selection with 3’ UTR design, this work redefines the 3’ UTR as an active translational enhancer and provides a broadly applicable regulatory element for next-generation mRNA and gene-delivery therapeutics.

---

Messenger RNA (mRNA) and adeno-associated viral (AAV) vectors have emerged as leading platforms for therapeutic protein expression.(*1–5*) mRNA enables rapid yet transient translation without genomic integration,(*6*) whereas AAV supports sustained expression through episomal persistence. Despite clinical successes in vaccines and genetic disorders,(*7*) both modalities remain constrained by inefficient translation(*8*), resulting from mRNA instability,(*9–11*) suboptimal ribosome recruitment,(*12, 13*) and limited processing of AAV-derived transcripts.(*14, 15*) Enhancing translational efficiency is therefore key to realizing the full potential of these gene expression platforms.

mRNA consists of a 5’ cap, untranslated regions (UTRs), coding sequence (CDS), and poly(A) tail, with the UTRs playing central roles in transcript stability and translational control.(*16*) While extensive optimization of 5’ UTRs through structural modeling and machine learning has improved expression,(*17–26*) progress on the 3’ UTR has lagged.(*27*) Current strategies—such as eliminating miRNA-binding sites,(*28, 29*) introducing AU-rich or viral-derived elements,(*30, 31*) or applying in vitro evolution(*32*)—primarily stabilize mRNA rather than enhance translation. Because efficient UTR regulation is also critical for AAV-delivered transgenes, new and versatile 3’ UTR design strategies are urgently needed for both mRNA- and AAV-based therapeutics.

Endogenous RNA regulation provides a natural model for translational enhancement.(*33*) The antisense long non-coding RNA Uchl1os activates translation of Uchl1 through an inverted SINEB2 repeat that functions as a cis-acting enhancer, coupled with an antisense domain conferring target specificity(*34–36*). Synthetic systems such as RNAe and CRISPR–dCasRx–SINEB2 have leveraged this SINEB2 domain to boost endogenous protein production.(*37, 38*) Incorporating such translation-enhancing motifs into therapeutic transcripts could overcome inefficient translation across delivery platforms.

Here we report an engineered 3’ UTR module that markedly enhances translation. Building on the natural activity of SINEB2, we used intracellular evolution to derive an optimized variant, P51, and its truncated form P51t3, which act in cis to recruit ribosomes through the closed-loop mechanism. When inserted into the 3’ UTR, both elements substantially increased protein output across multiple mRNA and rAAV constructs. Systematic validation and mechanistic analysis identified their molecular target, establishing P51 as a versatile 3’ UTR enhancer for improving translational efficiency and expanding the mechanistic understanding of active ribosome recruitment by 3’ UTRs.

## Intracellular evolution of the SINEB2 Element for enhanced protein expression

The SL1 stem–loop structure of SINEB2 is essential for its translation-enhancing activity.(*39*) Based on this, a 7-nucleotide segment within SL1 was randomized to generate a mutational library (**Fig.1A**). Screening employed a CRISPR–dCasRx–SINEB2 system,(*38*) in which HEK293F cells were engineered to stably express dCasRx and selected as monoclonal lines. A lentiviral library encoding crRNA–SINEB2 variants and puromycin N-acetyltransferase (PuroR) driven by the UbC promoter was introduced. In the cytoplasm, each SINEB2 variant was guided by dCasRx to PuroR mRNA, with each cell expressing one of 16,384 sequences (**Fig.1B**). Cells were cultured under gradually increasing puromycin pressure, genomic DNA was collected at multiple passages, and variant enrichment was analyzed by Sanger and next-generation sequencing (NGS) (**Fig.1C**).

**Fig. 1.**
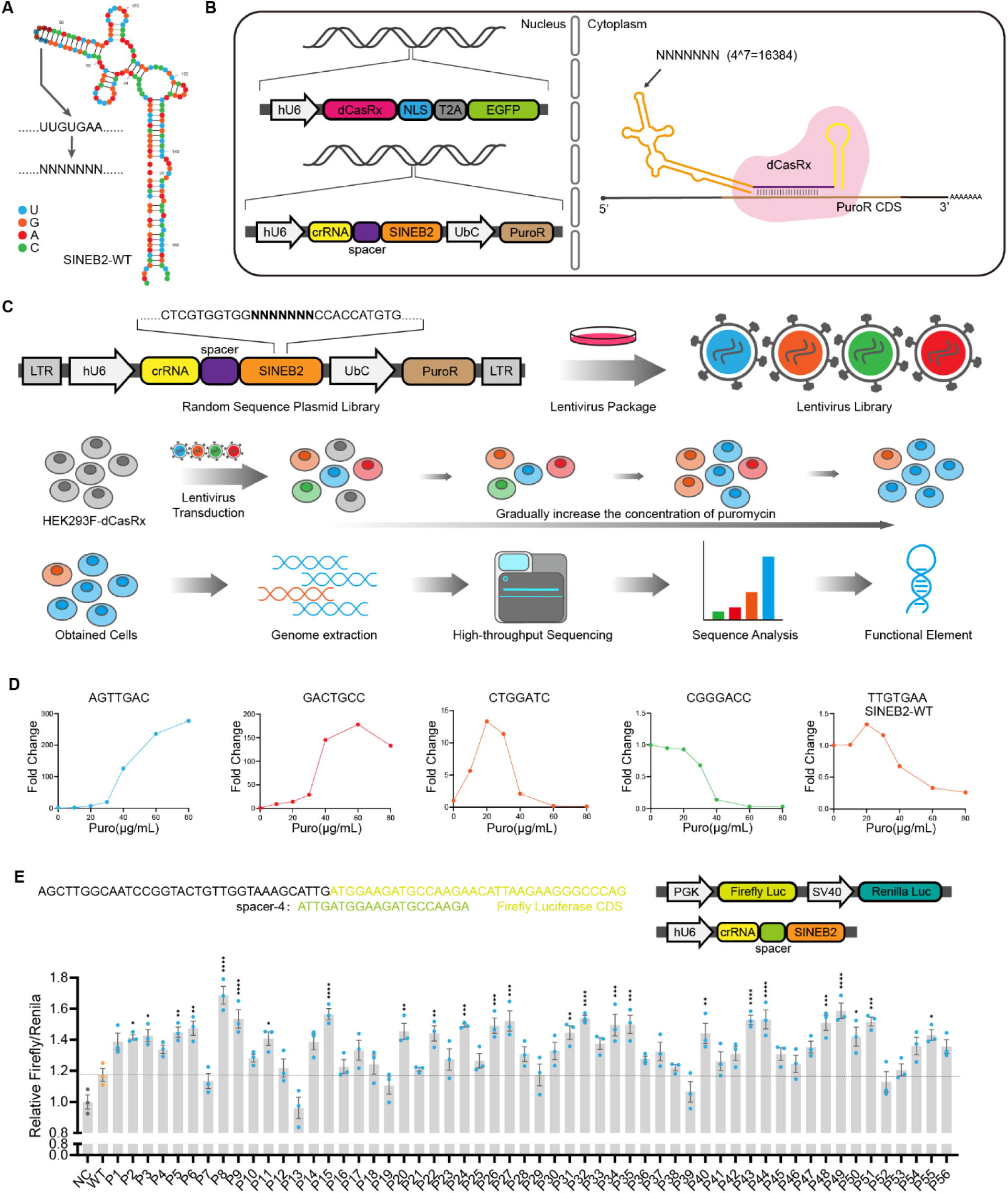
Intracellular evolution of SINEB2 elements. **(A),** The predicted secondary structure of SINEB2 and the sites to evolve. **(B)**, Schematic diagram of the expression of each component within a single cell during the screening process. Through two rounds of lentiviral transduction, the expression sequences of dCasRx, the crRNA-SINEB2 library, and PuroR were stably integrated into the cell genome. In the cytoplasm, dCasRx guided the crRNA-SINEB2 library to bind to the spacer-5 region of PuroR mRNA. Owing to MOI-controlled transduction, the majority of individual cells harbored only a single integrated sequence. The library encompassed a total of 16,384 distinct sequences. **(C)**, Schematic of the intracellular evolution and experimental procedure. Lentiviral plasmid library expressing crRNA-SINEB2 (4^7^=16384; U6 promoter) targeting near the PuroR start codon and PuroR (UbC promoter). Packaged lentivirus was transduced into HEK293F-dCasRx cells at MOI = 0.3. Cells underwent progressive puromycin selection (escalating from 0 to 80 μg/mL). Genomic DNA from endpoint cells was extracted and subjected to NGS of integrated crRNA. **(D),** The abundance changes of several representative sequences during the screening process. Fold Change is the proportion of a single sequence in the sequencing data divided by the proportion of that sequence in the initial library. **(E),** The function of the sequences obtained through screening was verified by the dual-luciferase reporter system, in HEK293F-dCasRx. The upper part indicates the plasmid structure and the localization of crRNA. NC (random sequence shuffled from SINEB2). Data are presented as mean ±SEM n =3. Two-tailed Student’s t-test. *P < 0.05, **P < 0.01, ***P < 0.001, ****P < 0.0001.

To establish the system, HEK293F lines expressing dCasRx with different numbers of nuclear localization signals (NLS) were created (**Fig. S1A**) and verified by EGFP fusion (**Fig. S1B**). Dual-luciferase reporter (DLR) assays identified the optimal SINEB2 target site in the NLS–dCasRx–NLS line. Spacer sequences at 8-nt intervals directed SINEB2-WT to various firefly luciferase (Fluc) regions, with site 4 giving maximal enhancement (**Fig. S1C–E**). The highest activation occurred with single-NLS dCasRx (**Fig. S1F**). To improve sensitivity and avoid translational saturation, transcription was driven by the weaker UbC promoter rather than EF1α (**Fig. S2A**). Spacer sites on PuroR mRNA were then tested, and spacer 5 yielded the strongest proliferative effect under puromycin selection, especially in dCasRx-NLS cells (**Fig. S2B–G**), and was thus used for screening.

Using the optimized monoclonal dCasRx-NLS line, spacer 5 was validated as the optimal gRNA target (**Fig. S3A,B**), and the crRNA–spacer–SINEB2 library was screened under stepwise puromycin (2–80 μg/mL). Colonies survived up to 60 μg/mL, while higher doses caused progressive cell loss (**Fig. S3C**). At 80 μg/mL, Sanger sequencing revealed shifts in base composition (**Fig. S3D**), and NGS across time points showed depletion of most variants but consistent enrichment of a few (**Fig. S3E**), identifying sequences with superior translation-promoting activity.

## Identification of the Highly Efficient Translation-Enhancing Element P51

NGS analysis identified four distinct enrichment patterns among library variants (**Fig. 1D**). Sustained enrichment sequences continuously increased, indicating strong translation-enhancing activity that supported growth under 80 μg/mL puromycin. Late decline sequences rose initially but fell at higher concentrations, suggesting moderate activity; early decline sequences dropped at lower concentrations, indicating weak activity; and continuous decline sequences decreased throughout, showing minimal effect. Wild-type SINEB2 (SINEB2-WT) exhibited an early rise followed by late decline, consistent with limited translation-enhancing capacity. From these profiles, 56 sustained enrichment candidates (P1–P56) were selected for validation.

Dual-luciferase assays in dCasRx-NLS cells confirmed that most P1–P56 variants outperformed SINEB2-WT (**Fig. 1E**). To test effects on endogenous genes, dCasRx-NLS was stably expressed in A375 cells (**Fig. S4A**), and gRNAs targeting the tumor suppressor PTEN were introduced. Cell viability assays showed that most variants suppressed proliferation more effectively than SINEB2-WT, consistent with enhanced PTEN translation (**Fig. S4B**). Western blotting further confirmed these findings, with P51 yielding the highest increase in PTEN protein (**Fig. S4C,D**). Secondary structure prediction revealed that while SINEB2-WT forms a stable stem–loop, P51’s 7-nucleotide mutation reconFigured the SL1 structure (**Fig. S4E**), likely underpinning its enhanced activity.

## P51 Enhances mRNA Translation via 3’ UTR Insertion

Having identified an RNA element that enhances translation, we next tested whether it could function when integrated into untranslated regions (UTRs), which would allow more convenient application. Previous studies showed that inserting an eIF4G aptamer into the 5’ UTR can promote translation.(*40, 41*) To assess whether SINEB2 acts similarly, SINEB2-WT was placed in the 5’ UTR of EGFP mRNA with optimized sequences around the AUG start codon. However, translation was reduced rather than enhanced (**Fig. S5A**), with inhibition varying by insertion site (**Fig. S5B**). This effect likely arose from the strong secondary structure of SINEB2 impeding ribosome scanning. Truncation to weaken the structure did not improve translation (**Fig. S5C**), indicating SINEB2 is unsuitable for 5’ UTR insertion (**Fig. S5D**).

In eukaryotes, translation efficiency is enhanced by the closed-loop model, where the 5’ cap–eIF4E and 3’ poly(A) – PABP complexes are bridged by eIF4G to facilitate ribosome recycling.(*42, 43*) Inspired by this, and by prior efforts tethering IRES elements to 3’ UTRs,(*44*) we inserted SINEB2-WT into the 3’ UTR following the RNAe strategy.(*37*) Complementary sequences to the 5’ UTR were introduced with varied positions and spacing (**Fig. 2A**). EGFP reporter assays showed modest enhancement **(Fig. 2B-D**), but removing the complementary sequence and adjusting spacer length increased translation by ∼1.5-fold (**Fig. 2E–G**), suggesting that duplex formation interfered with normal poly(A)–PABP–eIF4G interactions. Subsequent designs thus relied solely on closed-loop–mediated proximity.

**Fig. 2.**
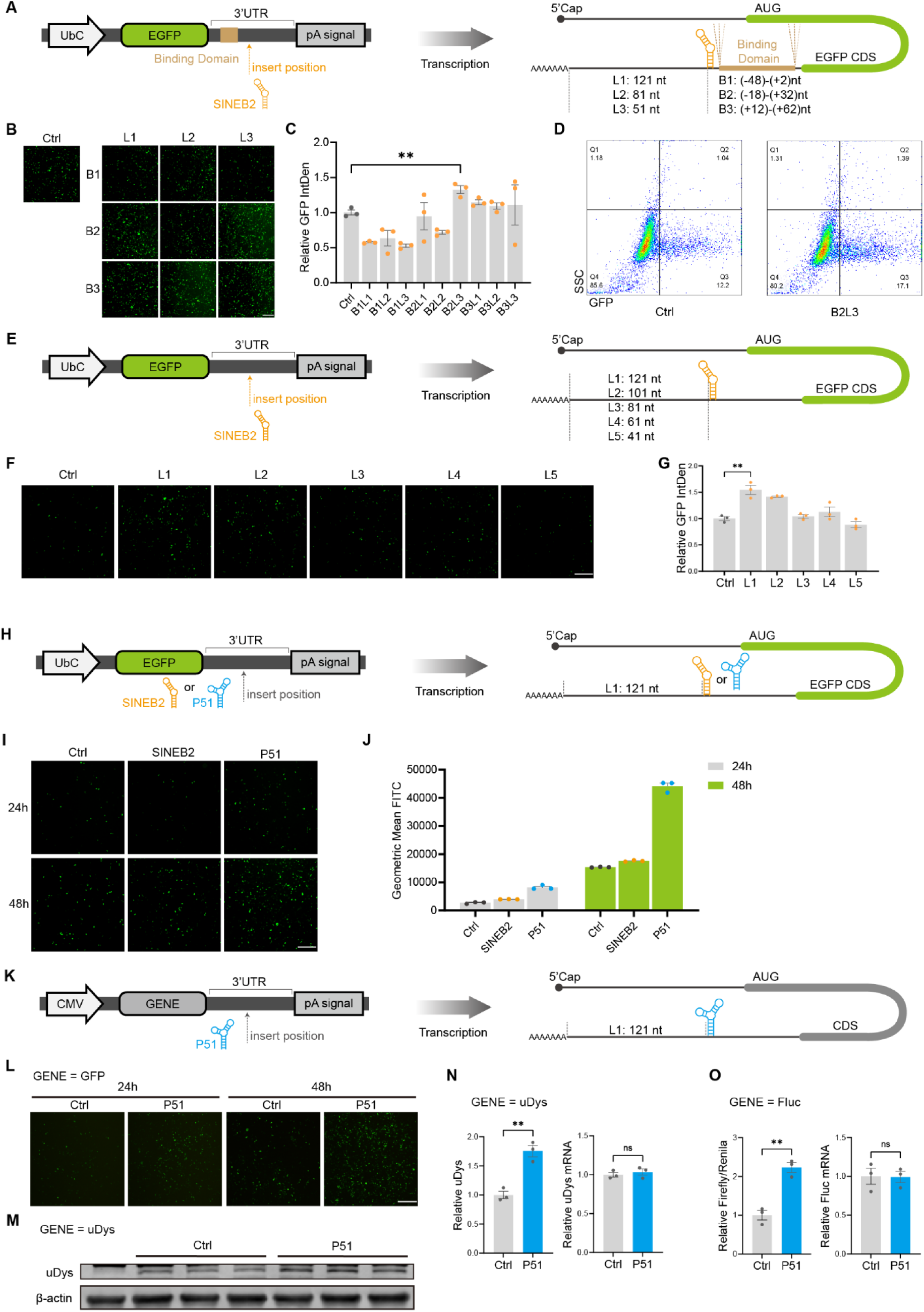
Engineering the 3’UTR with SINEB2 and P51 elements. **(A),** Schematic of Binding-SINEB2 elements inserted via different strategies in the 3’UTR. **(B),** The results of Binding-SINEB2 elements inserted via different strategies in the 3’UTR of UbC-EGFP was evaluated by fluorescence spectroscopy in HEK293T cells transfected with the respective engineered plasmids. Scale bar = 1000 µm. **(C),** Measurement of EGFP fluorescence enhancement mediated by Binding-SINEB2–engineered 3’UTRs. Data are normalized to the UbC-EGFP reporter 3’UTR lacking SINEB2 insertions (Ctrl) and presented as mean ± SEM (n = 3). **P < 0.01, two-tailed Student’s t-test. **(D),** Flow cytometry analysis showing the percentage of EGFP-positive cells mediated by Binding-SINEB2–engineered 3’UTR. **(E),** Schematic of SINEB2 elements without binding domain inserted at different sites in the 3’UTR. **(F),** The results of SINEB2 elements without binding domain inserted at different sites in the 3’UTR of UbC-EGFP was evaluated by fluorescence spectroscopy in HEK293T cells transfected with the respective engineered plasmids. Scale bar = 1000 µm. **(G),** Measurement of EGFP fluorescence enhancement mediated by 3’UTRs engineered with binding-domain-deficient SINEB2. Data are normalized to the UbC-EGFP reporter 3’UTR lacking SINEB2 insertions (Ctrl) and presented as mean ± SEM (n = 3). **P < 0.01, two-tailed Student’s t-test. **(H),** Schematic of SINEB2 element and selected P51 element inserted in the 3’UTR. **(I),** The results of SINEB2 element and selected P51 element inserted in the 3’ UTR of UbC-EGFP was evaluated by fluorescence spectroscopy in HEK293T cells transfected with the respective engineered plasmids. Scale bar = 1000 µm. **(J),** Flow cytometry analysis of fluorescence enhancement mediated by SINEB2 element and selected P51 element inserted in the 3’UTR.. **(K),** Schematic diagram of inserting the P51 element into the pCAG 3’UTR. There were 121 bases between the insertion site and polyA. After transcription, through the mRNA closed-loop translation model, P51 was spatially close to the 5’ region of mRNA. Data are normalized to the reporter 3’ UTR lacking P51 insertions (Ctrl). **(L),** P51 enhanced the translation of EGFP in HEK293T. Scale bar = 1000 µm. **(M),** P51 enhanced the translation of uDys. Western blot data from the HEK293T cells transfected with the plasmid at 48h post transfection. n=3. **(N),** The quantitative results of figure (d) and the RT-qPCR results of uDys mRNA. Data are presented as mean ±SEM n = 3. Two-tailed Student’s t-test. **P < 0.01, ns, not significant. **(O),** P51 enhanced the translation of Fluc in HEK293T. The results of the dual-luciferase reporter system and the RT-qPCR results of Fluc mRNA, from the HEK293T cells transfected with the plasmid at 48h post transfection. Data are presented as mean ±SEM n = 3. Two-tailed Student’s t-test. **P < 0.01, ns, not significant.

Replacing SINEB2-WT with the optimized P51 element (**Fig. 2H**) markedly improved translation, as shown by EGFP reporter and flow cytometry assays (**Fig. 2I,J**). Under the strong CMV promoter, P51 enhanced expression of multiple proteins in HEK293T cells, including EGFP (**Fig. 2K,L**), and also boosted production of the therapeutic micro-dystrophin (uDys)(*14*) without altering mRNA abundance (**Fig. 2M,N**). Similar results were observed with luciferase, where P51 increased protein output ∼2.1-fold while mRNA levels remained unchanged (**Fig. 2O**). Thus, 3’ UTR insertion of P51 significantly enhances translation efficiency without affecting stability, enabling a 2–3-fold improvement in protein expression with broad utility for gene therapy and recombinant protein production.

## The P51 Truncation Variant P51t3 Exhibits Broad Translation-Enhancing Activity

To test the universality of our optimization strategy, we evaluated P51 in different 3’ UTR contexts, using the Moderna mRNA-1273 3’ UTR as a model. Multiple insertion sites were designed at 20-nt intervals (**Fig.3A**, **Fig.S6A**), and reporter assays with EGFP and Fluc identified Site 2 as optimal (**Fig.3B,C**, **Fig. S6B**). Given the 167-nt length of P51, four truncated variants (P51t1–t4) were generated (**Fig.3D**). Structural predictions indicated that all retained P51’s basic features. When inserted at Site 2 (**Fig.3E**, **Fig. S6C**), P51t2 and P51t3 produced stronger enhancement than P51, achieving 4–6-fold translation increases relative to the blank UTR group (**Fig.3F,G**, **Fig. S6D**).

**Fig. 3.**
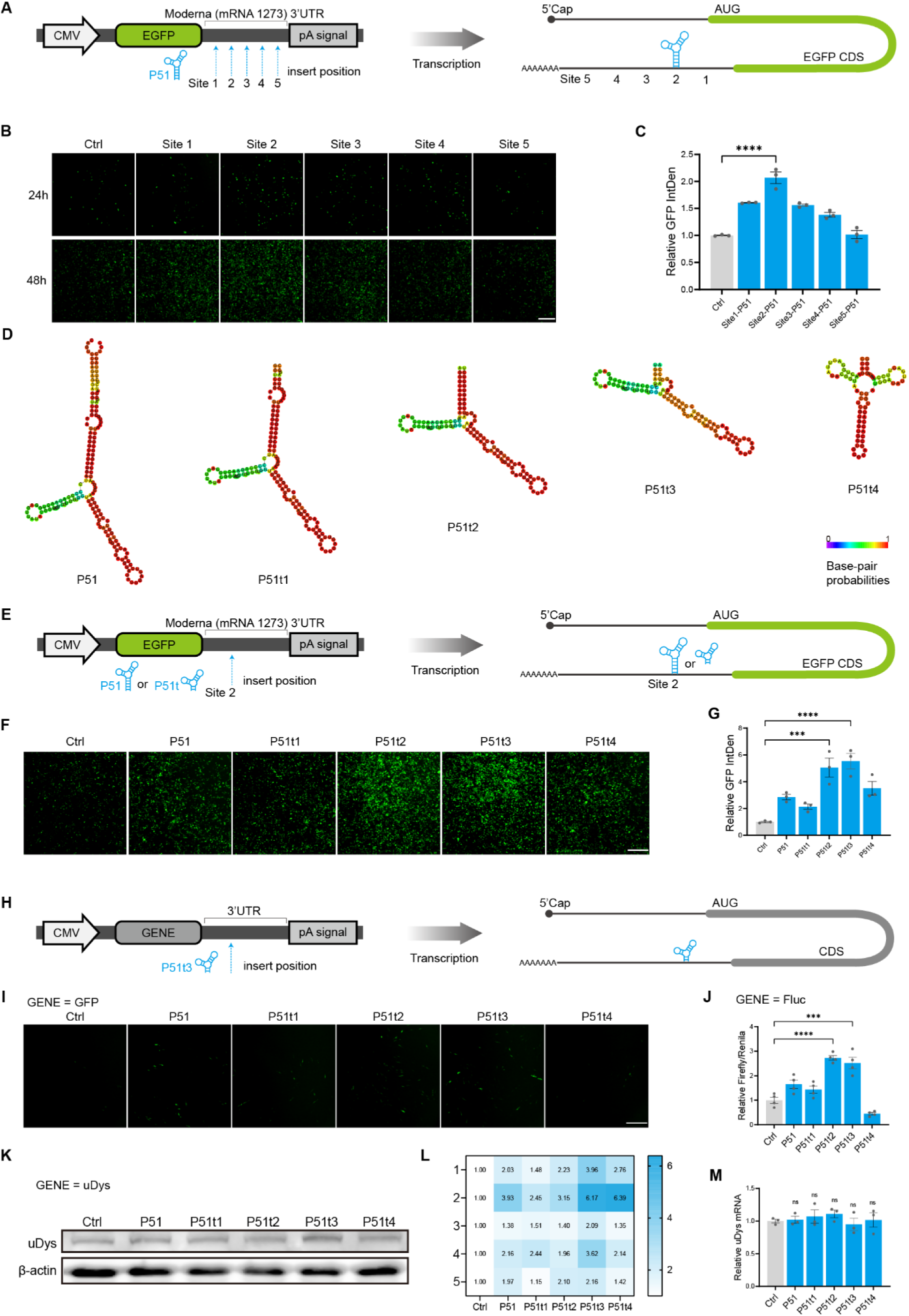
Validation and optimization of the P51 element in the 3’UTR. **(A),** Schematic of P51 elements inserted at five sites in the CMV-EGFP reporter 3’UTR. **(B),** The results of P51 elements inserted at five sites in the CMV-EGFP reporter 3’UTR (Moderna mRNA-1273 3’ UTR) was evaluated by fluorescence spectroscopy in HEK293T cells transfected with the respective engineered plasmids. Scale bar = 1000 µm. **(C),** Measurement of EGFP fluorescence enhancement mediated by P51 elements inserted at five sites in the CMV-EGFP reporter 3’UTR (Moderna mRNA-1273 3’UTR). Data are normalized to the CMV-EGFP reporter 3’UTR lacking P51 insertions (Ctrl) and presented as mean ± SEM (n = 3). **P < 0.01, two-tailed Student’s t-test. **(D),** Structures of the P51 element and its four truncations. RNA secondary structures were predicted using the RNAfold WebServer. **(E),** Schematic of P51 element and its four truncations inserted in the CMV-EGFP reporter 3’UTR. **(F),** The results of P51 element and its four truncations inserted in the CMV-EGFP reporter 3’UTR (Moderna mRNA-1273 3’UTR) was evaluated by fluorescence spectroscopy in HEK293T cells transfected with the respective engineered plasmids. Scale bar = 1000 µm. **(G),** Measurement of EGFP fluorescence enhancement mediated by P51 element and its four truncations inserted in the CMV-EGFP reporter 3’UTR (Moderna mRNA-1273 3’UTR). Data are normalized to the CMV-EGFP reporter 3’UTR lacking P51 or P51-truncation insertions (Ctrl) and presented as mean ± SEM (n = 3). ***P < 0.001, ****P < 0.0001, two-tailed Student’s t-test. **(H),** Schematic diagram of inserting the P51 element or its truncated forms into the pCAG 3’UTR. There were 121 bases between the insertion site and polyA. After transcription, through the mRNA closed-loop translation model, P51 or its truncated forms was spatially close to the 5’ region of mRNA. Data are normalized to the reporter 3’UTR lacking P51 or its truncated forms insertions (Ctrl). **(I),** P51t2 and P51t3 further enhanced the translation of EGFP in C2C12. Scale bar = 1000 µm. **(J),** P51t2 and P51t3 further enhanced the translation of Fluc in C2C12.The results of the dual-luciferase reporter system from the C2C12 cells transfected with the plasmid at 48h post transfection. Data are presented as mean ±SEM n = 4. Two-tailed Student’s t-test. ***P < 0.001, ****P < 0.0001. **(K),** P51t2 and P51t3 further enhanced the translation of uDys in C2C12. Western blot data from the C2C12 cells transfected with the plasmid at 48h post transfection. **(L),** Quantitative heat map of the experiment of figure (k). n=5. **(M),** The RT-qPCR results of uDys mRNA. Data are presented as mean ±SEM n = 4. Two-tailed Student’s t-test. ns, not significant.

We next validated truncated variants in additional contexts. In C2C12 cells, insertion of P51t2 or P51t3 into the pCAG 3’ UTR significantly increased expression of EGFP, Fluc, and micro-dystrophin (uDys), as confirmed by fluorescence, reporter assays, and Western blotting (**Fig.3H-L**). Enhancements ranged from 3–6 fold, while qPCR showed no changes in mRNA abundance (**Fig.3M**). Based on activity and compact size, P51t3 was selected for further studies. Testing in endogenous UTRs confirmed its versatility: insertion into B2M (six sites) and TMSB10 (nine sites) 3’ UTRs consistently enhanced FLUC translation(*45*) (**Fig. S6E-H**), independent of coding sequence. Enhancement followed a positional pattern, with one optimal insertion site per UTR and weaker effects at flanking positions, suggesting recruitment of regulatory proteins near the 5’ end.

To explore the mechanism, we analyzed potential miRNA interactions. Bioinformatic predictions identified several miRNA binding sites in full-length P51 (**Fig.S7A**),(*46*) one of which (miR-383-3p) was lost in truncated variants (**Fig.S7B,C**). As miR-383-3p is known to inhibit target mRNA expression,(*47*) elimination of this site may contribute to the superior activity of P51t2 and P51t3.

## The P51t3 Element Enhances Protein Expression in mRNA Vectors

After identifying P51t3 as the optimal element, we evaluated its performance in gene therapy–relevant in vitro transcribed (IVT) mRNA systems. Since P51t3 functions best near the 5’ end and IVT mRNAs differ in 5’ UTR length, insertion sites were re-screened. P51 was positioned at three sites spaced 20 nt apart (**Fig. 4A,F**), and reporter assays with Fluc and EGFP identified Site 3 as optimal (**Fig. 4B,G,H**). Among truncated variants, P51t3 again exhibited the strongest activity, consistent with plasmid results (**Fig. 4C,D**), whereas P51t2 performed worse than full-length P51 at Site 3, likely due to new miRNA-binding sites (**Fig. S7D**).

**Fig. 4.**
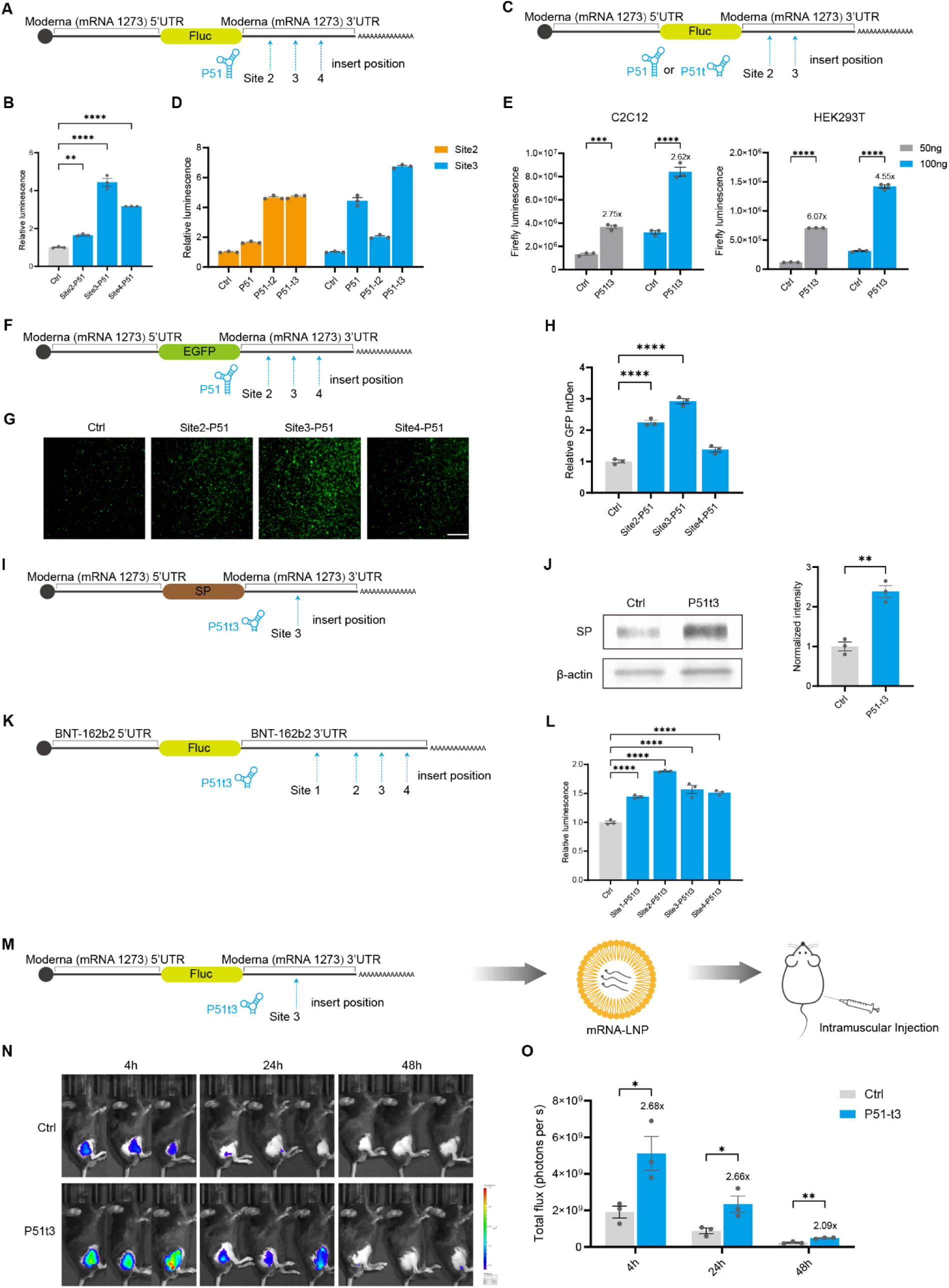
P51t3 element enhances the gene expression of the mRNA vector. **(A),** Schematic of P51 element inserted at three sites within the 3’UTR of IVT mRNA. **(B),** The results of P51 element inserted at three sites within the 3’UTR of IVT mRNA was evaluated by dual-luciferase assay in HEK293T cells transfected with the respective mRNAs. Data are normalized to the IVT mRNA 3’UTR lacking P51 insertions (Ctrl) and presented as mean ± SEM (n = 3). **P < 0.01, ****P < 0.0001, two-tailed Student’s t-test. **(C),** Schematic of P51 element or its truncations inserted at Site2 and Site3 within the 3’UTR of IVT mRNA. **(D),** The results of P51 element or its truncations inserted at Site2 and Site3 within the 3’UTR of IVT mRNA was evaluated by dual-luciferase assay in HEK293T cells transfected with the respective mRNAs. Data are normalized to the IVT mRNA 3’UTR lacking P51 insertions (Ctrl) and presented as mean ± SEM (n = 3). **(E),** The results of P51t3 element inserted at Site3 within the 3’UTR of IVT mRNA was evaluated by dual-luciferase assay in C2C12 and HEK293T cells transfected with 50 ng (gray) or 100 ng (blue) of the respective mRNAs. Data are normalized to the IVT mRNA 3’UTR lacking P51t3 insertions (Ctrl) and presented as mean ± SEM (n = 3). ***P < 0.001, ****P < 0.0001, two-tailed Student’s t-test. **(F),** Schematic of P51 elements inserted at three sites within the 3’UTR (Moderna mRNA-1273 3’UTR) of IVT EGFP mRNA. **(G), (H),** The results of P51 elements inserted at three sites within the 3’UTR (Moderna mRNA-1273 3’UTR) of IVT EGFP mRNA was evaluated by fluorescence spectroscopy in HEK293T cells transfected with the respective mRNAs. Scale bar = 1000 µm. EGFP data are normalized to the IVT mRNA 3’UTR lacking P51t3 insertions (Ctrl) and presented as mean ± SEM (n = 3). ****P < 0.0001, two-tailed Student’s t-test. **(I),** Schematic of P51t3 element inserted at Site3 within the 3’UTR of IVT Moderna mRNA-1273. **(J),** The results of P51t3 element inserted at Site3 within the 3’UTR of IVT Moderna mRNA-1273 was evaluated by western blot assay in HEK293T cells transfected with the mRNAs. Data are normalized to the IVT mRNA 3’UTR lacking P51t3 insertions (Ctrl) and presented as mean ± SEM (n = 3). **P < 0.01, two-tailed Student’s t-test. **(K),** Schematic of P51t3 elements inserted at four sites within the 3’UTR (BioNTech/Pfizer BNT-162b2 3’UTR) of IVT Fluc mRNA. **(L),** The results of P51t3 elements inserted at four sites within the 3’UTR (BioNTech/Pfizer BNT-162b2 3’UTR) of IVT Fluc mRNA was evaluated by dual-luciferase assay in HEK293T cells transfected with the respective mRNAs. Data are normalized to the IVT mRNA 3’UTR lacking P51t3 insertions (Ctrl) and presented as mean ± SEM (n = 3). ****P < 0.0001, two-tailed Student’s t-test. **(M),** Schematic of in vivo luminescence quantification. Fluc mRNAs were formulated in LNPs and equal molar quantities of mRNA-LNP complexes were administered through IM injection to male 6-8 weeks C57BL/6J mice. **(N), (O),** Exemplary in vivo luminescence images of mice treated with Fluc mRNA at 4, 24 and 48 h post mRNA–LNP complex administration. Color scale of the heatmaps, radiance (photons s−1 cm−2 sr−1). Luminescence was measured by integration of total flux for each mouse. Data are normalized to the mice of Ctrl group (IVT mRNA 3’UTR lacking P51t3 insertions-LNP-treated) and presented as mean ± SEM (n = 3). *P < 0.05, **P < 0.01, two-tailed Student’s t-test.

P51t3 performance was then tested under varied conditions. Across human and murine cells and multiple mRNA doses, IVT mRNAs carrying P51t3 achieved 2.5–6-fold higher protein output (**Fig. 4E**). In clinically relevant models, insertion of P51t3 into an mRNA containing Moderna mRNA-1273 UTRs increased spike protein expression 2–3-fold in C2C12 cells (**Fig. 4I,J**).48 Similarly, in constructs mimicking BioNTech BNT162b2 UTRs, P51t3 boosted Fluc translation nearly 2-fold (**Fig. 4K,L**).

In vivo, lipid nanoparticle (LNP)–encapsulated mRNAs were injected intramuscularly into mice. Bioluminescence imaging at 4, 24, and 48 hours post-injection showed a consistent 2–3-fold increase in expression with P51t3 (**Fig. 4M–O**). Collectively, these results establish P51t3 as a versatile 3’ UTR regulatory module that substantially enhances translation efficiency across diverse sequence contexts and delivery platforms, underscoring its potential for gene therapy and mRNA-based therapeutics.

## The P51t3 Element Enhances Protein Expression in rAAV Vectors

Recombinant adeno-associated virus (rAAV) is a leading vector for gene therapy owing to its safety, durability, and tissue specificity.(*4*) By replacing the wild-type gene with a therapeutic payload, rAAV enables targeted, long-term expression. After entry, the single-stranded DNA genome circularizes, undergoes second-strand synthesis, and is transcribed into mRNA to produce the therapeutic protein(*4*). However, suboptimal expression often limits efficacy, whereas dose escalation risks hepatotoxicity, immune activation, and off-target effects.(*48*)

To test whether P51t3 enhances rAAV expression, we constructed an rAAV plasmid containing the validated pCAG 3’ UTR and packaged it for testing in C2C12 cells (**Fig. S8A**). Initial assays showed no enhancement by P51 or its truncations (P51t2, P51t3) (**Fig. S8B,C**), likely due to proximity of the element to the 3’ inverted terminal repeat (ITR), which may hinder second-strand synthesis or circularization. Inserting a random spacer between the poly(A) signal and ITR improved expression (**Fig. S8D**). Combining this spacer with truncation optimization (**Fig. S8E**) enabled P51t3 to increase long-term protein expression 4–5 fold in C2C12 cells (**Fig. S8F**). Using micro-dystrophin (uDys) as a therapeutic model, the P51t3–In2 modification further enhanced expression (**Fig. S8G,H**).

In vivo, rAAV–Fluc vectors were injected intramuscularly into mice (**Fig. S8I**). Bioluminescence imaging at days 4, 8, 12, and 60 (**Fig. S8J**) showed that P51t3–In2 achieved ∼6-fold higher expression compared to control vectors (**Fig. S8K**). Together, these data demonstrate that P51t3 significantly enhances translation across plasmid, IVT mRNA, and rAAV systems, highlighting its potential as a broadly applicable regulatory element for therapeutic development.

## Ribosome Recruitment by SINEB2-WT and P51 through Interaction with RPL39

Two mechanisms have been proposed for SINEB2-mediated translation enhancement: recruitment of initiation factors or direct binding to the large ribosomal subunit.(*49–51*) To test the first, Fluc reporters with varying 5’ UTR lengths were analyzed in the CRISPR–dCasRx–SINEB2 system (**Fig. S9A,B**). If SINEB2 recruited initiation factors, the optimal spacer site would shift with UTR length, but it remained constant (**Fig. S9C**), suggesting initiation factor recruitment is not dominant.

We next identified protein partners of SINEB2-WT and P51. RNA–protein interaction screening in A375–dCasRx-NLS cells followed by anti-dCasRx immunoprecipitation and mass spectrometry (**Fig. 5A**) revealed enrichment of ribosomal protein RPL39—a 60S subunit component(*52*) —by both SINEB2-WT and P51 (**Fig. 5B,C**), with no change in initiation factor abundance. CoIP–WB confirmed the interaction, showing stronger binding for P51 (**Fig. 5D**). siRNA-mediated RPL39 knockdown, verified by qPCR and Western blotting (**Fig. 5E,F**), markedly reduced translation enhancement by SINEB2-WT, P51, P51t2, and P51t3 (**Fig. 5G,H**). Bio-layer interferometry demonstrated that P51 binds RPL39 with high affinity (KD = 21.1 pM), confirming direct interaction (**Fig. 5I**).

**Fig. 5.**
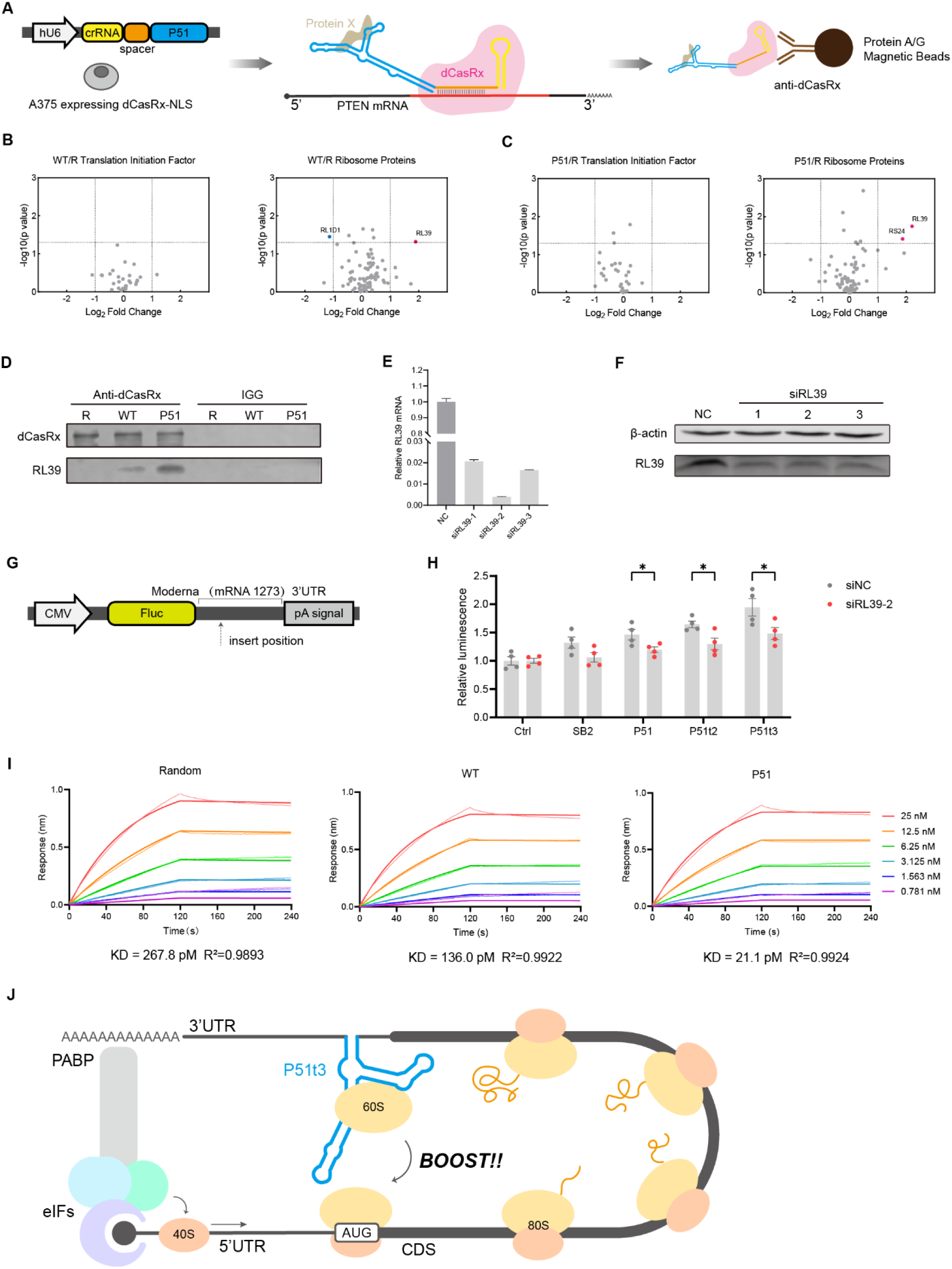
P51t3 element enhances mRNA translation by binding to RL39 and recruits the ribosomal 60S subunit. **(A),** Schematic diagram of target protein identification. (i) Transfect the crRNA-WT/P51 expression plasmid into A375-dCasRx cells. (ii) Fix the cells through formaldehyde. dCasRx, crRNA-WT/P51 and the target protein form a complex. (iii) Lysate the cells. dCasRx antibody was bound to Protein A/G magnetic beads. The antibody bound to the dCasRx protein complex. Mass spectrometry detection was carried out after magnetic separation. **(B), (C),** The result of protein mass spectrometry. The translation initiation factors and ribosome-related proteins were statistically analyzed respectively. WT and P51 were respectively compared with random sequence. Data are presented as mean ±SEM n =3. Two-tailed Student’s t-test. **(D),** The co-immunoprecipitation results of dCasRx and RL39. Random sequence, WT and P51 were compared. **(E),** The results of RT-qPCR that knocked down RL39. Data are presented as mean ±SEM n = 3. **(F),** The results of Western blot that knocked down RL39. **(G),** Schematic diagram of the plasmid for functional verification of RL39. Insert WT, P51, P51t2 and P51t3 at the insertion sites respectively. **(H),** Quantitative results of the luciferase reporter system. 48 hours after transfection with siRNA, transfect the plasmid, and then conduct the test 48 hours later. Data are presented as mean ±SEM n = 4. **(I),** BLI assay to determine the KD value of WT and P51. **(J),** Model for the action mechanism of P51t3 element in 3’UTR.

Consistent with Sharma et al.,(*50*) structural analysis suggested that SINEB2-WT may also engage rRNA via base pairing, and in P51 this region displays reduced base-pairing energy and increased positional entropy (**Fig. S9D,E**), implying enhanced rRNA accessibility.

Together, these findings support a model in which SINEB2-WT recruits the 60S subunit via RPL39 and rRNA interactions, while P51 strengthens these associations to accelerate translation initiation. In this model (**Fig. 5J**), mRNA circularization through cap–eIF4E and PABP–poly(A) interactions positions the 3’ UTR element near the start codon, allowing P51t3 to pre-recruit 60S subunits via RPL39–rRNA binding. When the 40S subunit completes scanning and reaches AUG, rapid 80S assembly ensues, thereby boosting translational efficiency.

## Discussion

mRNA therapeutics have transformed vaccine and protein replacement strategies but remain limited by inefficient translation.(*6*) Here, we developed a translation-enhancing RNA element, P51, and its optimized truncation variant P51t3, through intracellular directed evolution of engineered 3’ UTRs. P51t3 consistently improved protein output across multiple systems, establishing a universal regulatory module and providing both conceptual and technical advances in RNA element design.

Most optimization efforts target the 5’ UTR, codon usage, or nucleotide modifications, whereas 3’ UTR engineering has largely focused on transcript stability by removing miRNA sites or adding stabilizing motifs.(*2, 16, 27*) Our study introduces a distinct paradigm—embedding a ribosome-recruiting element within the 3’ UTR to directly boost translation. Using a CRISPR–dCasRx-based intracellular evolution platform, we recreated the native translation environment and applied selective pressure to identify high-performing variants. This strategy not only yielded potent enhancers such as P51 and P51t3 but also demonstrated the broader promise of intracellular evolution for discovering functional noncoding RNA elements. Scaling to larger libraries or integrating computational design could further improve translation efficiency.

A key finding is that SINEB2 derivatives are ineffective when placed in the 5’ UTR, likely because their stable secondary structures interfere with ribosomal scanning. In contrast, insertion into the 3’ UTR leverages the natural closed-loop translation mechanism, positioning the element near the start codon without hindering initiation. This demonstrates that the 3’ UTR, traditionally viewed as a region controlling stability and decay, can also be engineered as an active regulator of translation initiation.

Mechanistic analyses revealed that SINEB2 derivatives are ineffective in the 5’ UTR, likely due to secondary structures impeding ribosomal scanning. In contrast, 3’ UTR insertion exploits the natural closed-loop translation mechanism, positioning the element near the start codon without blocking initiation. This finding highlights that the 3’ UTR—traditionally viewed as a region for stability and decay regulation—can also be engineered as an active driver of translation initiation.

Further studies showed that P51 and P51t3 enhance translation not through canonical initiation factors but by directly interacting with ribosomal protein RPL39, a component of the 60S subunit, potentially aided by complementary rRNA interactions. Binding assays confirmed high-affinity binding, and RPL39 knockdown impaired the activity of SINEB2 and its optimized derivatives. These results expand the mechanistic scope of 3’ UTR biology, revealing that engineered elements can actively recruit ribosomes and accelerate 80S assembly at the start codon.

Functionally, P51t3 proved universal across diverse 3’ UTRs, coding sequences, promoters, and delivery systems. It enhanced expression in both human and murine cells, under weak or strong promoters, and in genes ranging from reporters to therapeutics such as micro-dystrophin. P51t3 remained active in clinically validated vaccine backbones, including Moderna mRNA-1273 and BioNTech BNT162b2, underscoring compatibility with real-world applications. In rAAV vectors, we identified a positional constraint near the ITR, which was resolved by inserting a spacer sequence to restore high expression. This reveals an unappreciated design variable in rAAV architecture and offers a generalizable optimization strategy. By achieving higher expression per construct, P51t3 could reduce vector or lipid nanoparticle (LNP) doses required for therapeutic efficacy, improving both safety and cost.

Together, these findings establish P51t3 as a universal 3’ UTR module that broadens the functional repertoire of noncoding regions and provides a practical solution to one of the central bottlenecks in mRNA therapeutics. Integrating intracellular directed evolution with rational UTR engineering creates a powerful framework for discovering new RNA elements. Future work may expand mutational landscapes, incorporate predictive algorithms, and combine P51t3 with other optimization strategies.

## Acknowledgments

The authors extend our thanks to the staff at the Peking University Medical and Health Analysis Center and the State Key Laboratory of Natural and Biomimetic Drugs for their invaluable assistance with instrumental analysis. This work is financially supported by the National Key R & D Program of China (Grant No. 2023YFF1205902, 2022YFA1304501), the National Natural Science Foundation of China (Grant No. 22227805, 22374004), Excellent Young Scientists Fund Program (Overseas) and Peking University. Correspondence and requests for materials should be sent to Dr. Liqin Zhang.

## Author contributions

Conceptualization: X.L., Q.Z., and L.Z.

Methodology: X.L., Q.Z., J.W., and Z.Z.

Investigation: X.L.

Visualization: X.L. and Q.Z.

Project administration: L.Z.

Supervision: L.Z.

Writing – original draft: X.L., Q.Z.

Writing – review & editing: L.Z.

## Data and materials availability

## Lead contact

Requests for further information and resources should be directed to and will be fulfilled by the lead contact, Liqin Zhang..

## Materials availability

This study did not generate new unique materials.

Plasmids generated in this study were listed in Table S2.

## Data and code availability

All data supporting the findings of this study are included within the article and its supplemental information and are also available from the authors upon reasonable request.

## Supplementary Materials

## Materials and Methods

### Cell Culture Condition

HEK-293T, C2C12 and A375 cells were cultivated in DMEM supplemented with 10% (v/v) fetal bovine serum (FBS), 10 μg/mL streptomycin, and 100 units/mL penicillin at 37°C with 5% CO2 in a humidified incubator. During adherent culture, HEK-293F was cultivated in MEM/EBSS supplemented with 10% (v/v) fetal bovine serum (FBS), 1% (v/v) non-essential amino acid (100x NEAA), 10 μg/mL streptomycin, and 100 units/mL penicillin at 37°C with 5% CO2 in a humidified incubator. During suspension culture, HEK-293F was cultured in SMM 293-TII Expression Medium, 10 μg/mL streptomycin, and 100 units/mL penicillin at 37°C with 5% CO2 in a humidified incubator.

### Construction of lentiviral plasmid libraries

First, the pLV-U6-PuroR-spacer-5-BsaI-UbC-PuroR plasmid containing restriction enzyme sites was constructed using sequence synthesis and GoldenGate assembly. Next, primers containing BsaI sites and a 7-nucleotide random sequence were synthesized(BsaI-SINEB2-N7-F : atgcGGTCTCGGATCCCCCAGAACTGGAGTTATACGGTAACCTCGTGGTGGNNNNNNNCCACCATGTGGATGGATATTGAGTTCCAAACACTGGTCCTGTGCAAG; BsaI-SINEB2-R : atgcGGTCTCGTCGACgaattCAAAAAAAGGAGCTAAAGAGATGGCTCAGCACTTAAGAGCACTGGATGCTCTTGCACAGGACCAGTGTTTGGAACTCAATAtcc). The library insert fragments were then generated by overlap extension PCR. Following digestion, the products were ligated using T4 DNA ligase. The ligation mixture was transformed into Trelief® 5α chemically competent cells. After amplification, the plasmids were extracted.

### Lentivirus package

To generate lentivirus, the lentivirus plasmid and packaging plasmid were transiently transfected into HEK293T cells. After transfection for 48 h and 72 h, the supernatant was filtered through a 0.45 μm filter. The 5X virus concentrate (PEG8000) was concentrated overnight at 4 ℃, and the virus particles were collected by centrifugation at 4000 × g for 30 minutes at 4 ℃. Resuspend the virus particles in cold medium and store at -80℃ or transduct the cells. Titer detection was performed using the Human Immunodeficiency Virus type 1 (HIV-1) p24 ELISA Kit before use.

### Plasmid construction

For construction of the 5’UTR-engineered UBC-EGFP plasmid, the SINEB2 element and its mutants were synthesized and annealed to form double-stranded inserts. These were PCR-extended with UBC 5’UTR segments and cloned into the EcoRI/KpnI sites of the parental vector.

For construction of the 3’UTR-engineered UBC-EGFP plasmid, 3’UTR sequences containing either the SINEB2 element or the P51 element were obtained by inserting them into the NotI/HindIII restriction sites of the parental UBC-EGFP vector.

For construction of the Moderna mRNA-1273 3’UTR-engineered CMV-EGFP/Fluc/SpikeProtein plasmid, the 5’UTR and 3’UTR were first replaced with the UTR sequences from Moderna mRNA-1273 to generate a control plasmid. Subsequently, the P51 element or P51 truncations were inserted into multiple specific sites within the 3’UTR using the GoldenGate method to complete the plasmid construction.

For construction of the BioNTech/Pfizer BNT-162b2 3’UTR-engineered CMV-Fluc plasmid, the 5’UTR and 3’UTR were first replaced with the UTR sequences from BioNTech/Pfizer BNT-162b2 to generate a control plasmid. Subsequently, the P51t3 element was inserted into multiple specific sites within the 3’UTR using the GoldenGate method to complete the plasmid construction.

To construct the rAAV plasmid, the CDS sequence was first replaced. Subsequently, the P51t3 element was inserted into a specific site within the 3’UTR using the homologous recombination method to complete the construction of the plasmid. Homologous recombination was also adopted when inserting spacer sequences.

### Transfection

For plasmid DNA transfection, HEK293T, C2C12 and A375 cells were seeded in 96-well plates or 6-well plates and allowed to grow overnight to reach 70-80% confluency. Transfections were performed using PEI according to the manufacturer’s instructions. Unless otherwise specified, for transfection in 96-well plates, 10-100 ng of plasmid DNA was used per well, and for 6-well plates, 1-2 µg of plasmid DNA was transfected per well.

For mRNA transfection, HEK293T or C2C12 cells were seeded in 96-well plates or 6-well plates and allowed to grow overnight to reach 70-80% confluency. Transfections were performed using NanoTrans™ Transfection Reagent Plus (CYTOCH, CT0005) according to the manufacturer’s instructions. For 96-well plates, 50 ng or 100 ng of mRNA was transfected per well. In the case of 6-well plates, 1 µg of mRNA was used per well.

### Recombinant AAV production

HEK293 cells were plated in 15 cm dishes at a density of 2 x107 cells/dish. The next day, each plate was transfected with pHelper plasmid, pAAV-RC9 plasmid, and ITR-containing plasmid using PEI. Recombinant virus was harvested from the cells and media, and purified by ultracentrifuge using a iodixanol gradient as previously described. AAV titers were quantified by qPCR. Primer and probe sequences for qPCR are listed in table S2.

### In vitro transcribed RNA preparation

For IVT of mRNA, DNA templates containing the T7 RNAP promoter were derived from suitable plasmids and were amplified by PCR before transcription. The template for mRNA transcription included a poly(A) tail (30A + GCATATGACT + 70A) at its 3’ end, which was added using a lengthy reverse primer during PCR. For a 30 µL reaction, 2 µg of purified template was incubated with components from the T7 Co-transcription RNA Synthesis Kit (CYNBIO, C3050) at 37 °C overnight. On the subsequent day, the RNA was purified using RNA purification magnetic beads (Transgene, EC501).

### FACS analysis to measure EGFP protein expression

Cells were trypsinized, washed once in complete media, then resuspended in PBS. At least 10,000 cells were measured on CytoFLEX flow cytometer and FACS data were analyzed using FlowJo software. GFP protein expression corresponds to EGFP mean fluorescence intensity (MFI).

### Luciferase assay

For the dual-luciferase reporter assay, HEK293T cells were seeded in 96-well plates and allowed to grow overnight to 70-80% confluency. The cells were co-transfected with 10 ng of Renilla plasmid and 50 ng of Firefly plasmid per well using PEI (Yeasen, 40816ES02). Forty-eight hours post-transfection, the culture medium was aspirated, and the cells were lysed with the lysis buffer provided in the Dual-luciferase Reporter Gene Detection Kit (Beyotime, RG028) for 15 minutes. The lysate was then transferred to a 96-well opaque plate. The luminescence was measured using a microplate luminometer (Centro, LB960) equipped with the capability to aspirate and inject luciferase substrate solution.

For the Firefly luciferase assay, HEK293T or C2C12 cells were seeded in 96-well plates and allowed to grow overnight to 70-80% confluency. The cells were transfected with 50ng or 100 ng of Firefly mRNA per well using NanoTrans™ Transfection Reagent Plus (CYTOCH, CT0005). Six hours post-transfection, the culture medium was aspirated, and the cells were lysed with the lysis buffer provided in the Dual-luciferase Reporter Gene Detection Kit (Beyotime, RG028) for 15 minutes. The lysate was transferred to a 96-well opaque plate, and the luminescence was measured using a microplate luminometer (Centro, LB960) equipped with the capability to aspirate and inject luciferase substrate solution.

### Western blotting

The treated cells were washed with PBS and lysed in RIPA buffer (Epizyme, PC103) supplemented with protease inhibitors and phosphatase inhibitors (Beyotime, P1005). After 15 min incubation at 4 °C, the lysates were centrifuged to remove debris. Total protein concentration was measured by BCA assay (Epizyme, ZJ102). 10 μg to 35 μg total protein was boiled in protein loading buffer (Epizyme, LT101) for 5 min at 95 °C and loaded onto 7.5–12.5 % SDS-PAGE gel according to target protein size. The total protein amount loaded was confirmed to be within linear range of detection for each antibody to detect each target protein. After stacking at 80 V, the gel was run at 120 V until the dye front reached the bottom. The proteins were transferred onto a methanol activated PVDF membrane (pore size 0.22 µm or 0.45 µm; Millipore) using wet transfer system. Membranes were blocked with rapid blocking buffer for 15 min at room temperature, incubated with primary antibody at 4 °C overnight, and then washed with TBST buffer (3 × 5 min), followed by corresponding fluorescent secondary antibody incubation 1 h at room temperature. The loading control β-Actin was visualized using 1:10000 fluorescent anti-β-Actin antibody. Membranes were imaged on Odyssey® DLx Dual-color Infrared Laser Imaging System.

### Co-inmunoprecipitation

First, the A375-dCasRx-NLS cell line was transfected with plasmid vectors expressing the corresponding crRNA sequences. 48 hours post-transfection, the cells were harvested, fixed with formaldehyde, and quenched with glycine. The cells were then lysed using Cell Lysis Buffer to extract total protein. Subsequently, the total protein lysate was incubated overnight at 4°C with rotation with Protein A/G Magnetic Beads that had been pre-bound with anti-dCasRx antibody or control IgG. After thorough washing, the beads were eluted using SDS loading buffer under denaturing conditions. The eluted samples were analyzed by both mass spectrometry and Western blot.

### Mice

All animal protocols were approved by the Animal Care and Use Committee of Peking University. C57BL/6J mice, 19-20 g, were maintained under standard laboratory conditions (a strict 12-hour light/dark cycle, a controlled temperature of 23 ± 2°C and a humidity at 40–70%). All mice were randomly assigned to experimental groups, and no mice were excluded from analysis.

### In vivo delivery and imaging of FLuc

After assembly into lipid nanoparticles, equal molar amounts of control Firefly mRNA and Firefly mRNA with engineered P51t3 in the 3’UTR were administered via intramuscular injection. Similarly, after AAV packaging and purification, intramuscular injection was administered to mice at a dose of 10¹¹ vg per site. The in vivo activity of Firefly was measured using the IVIS Lumina Series III. Freshly prepared D-Luciferin Potassium Salt substrate was dissolved in PBS. Mice were anesthetized and imaged under default settings. Luminescent activity was quantified using Living Image.

### Bio-Layer Interferometry

The binding affinity of SINEB2 and P51 to the RL39 was assessed using biolayer interferometry (BLI). The binding experiments were performed in black 96-well plates, with each well containing 200 μL of BLI buffer (50 mM Tris-HCl, 5 mM MgCl2, 1 mM CaCl2, 0.1% Tween 20). NTA sensors were used for binding the RL39. The sensors were prehydrated with BLI buffer for a minimum of 10 minutes. To generate analyte samples, aptamer-based degraders were dissolved in BLI buffer and then diluted to five different concentrations. The experiments followed a five-step sequential assay: Baseline (60 s); RL39 loading (120 s); Baseline (60 s); Association (200 s); Dissociation (200 s). The data were primarily processed with Octet BLI Analysis 12.2, exported as a dataframe, and the illustration was recreated using GraphPad.

### Quantification and Statistical Analysis

Details of exact statistical analysis, packages, tests, and other procedures used can be found in the main text, Figure legends, and STAR Methods.

---

**Fig. S1.**
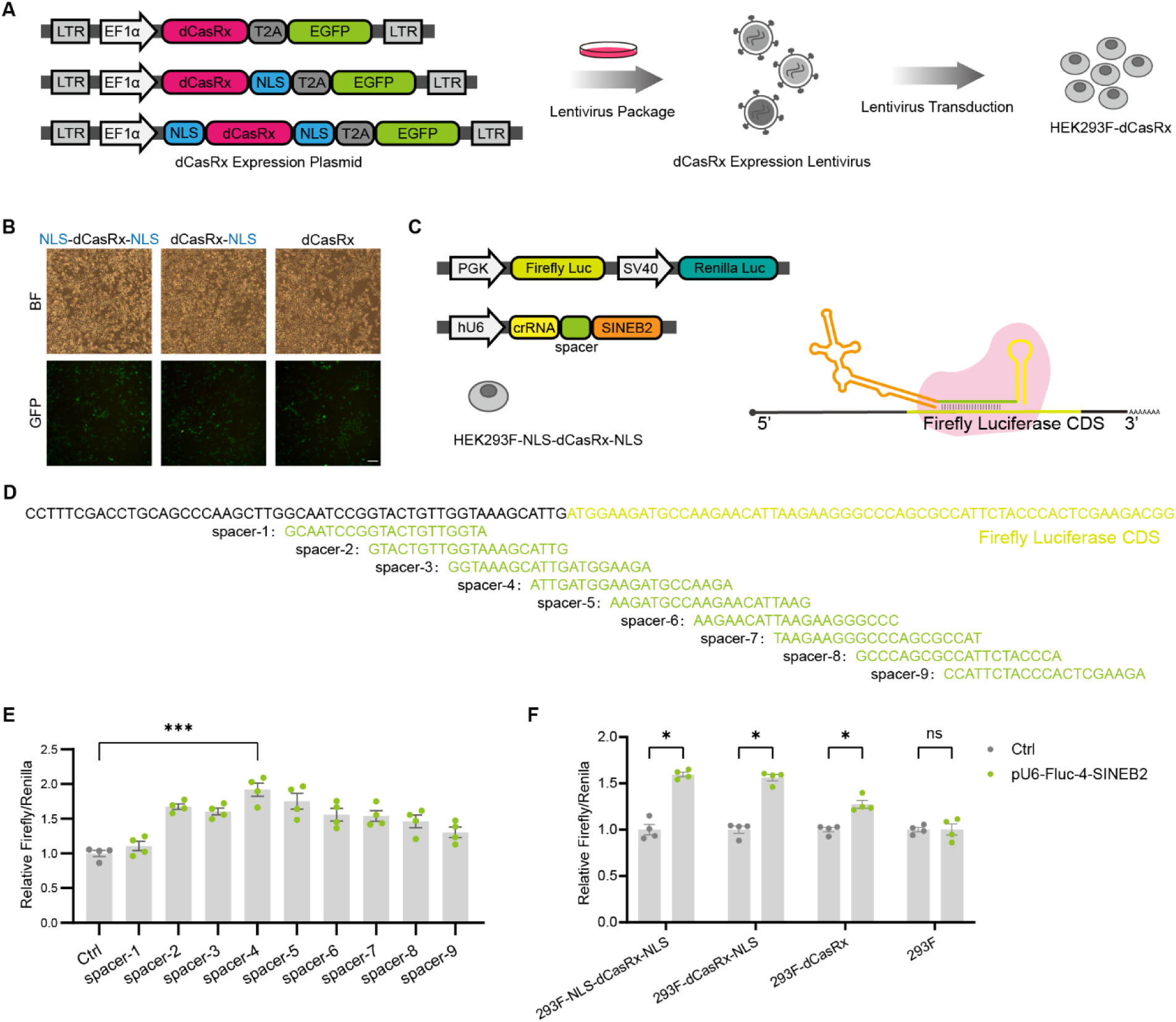
dCasRx cell line used for intracellular evolution, related to Fig 1. **(A),** Schematic diagram of establishing HEK293F cell Lines expressing dCasRx with various nuclear localization signals via lentiviral transduction. **(B),** The expression of EGFP co-translated with dCasRx via T2A. Scale bar = 200 µm. **(C),** The function of SINEB2 was verified by a dual-luciferase reporter system. crRNA-SINEB2 was located on the mRNA of firefly luciferase via dCasRx. **(D),** Different spacer sites of firefly luciferase mRNA. **(E),**The dual luciferase assay verified the upregulation of firefly translation by SINEB2 at different spacer sites. Ctrl (pcDNA3.1). Data are presented as mean ±SEM n =4. Two-tailed Student’s t-test. ***P < 0.001. **(F),** The dual luciferase assay verified the upregulation of firefly translation by different NLS at the same spacer site. Ctrl (pcDNA3.1). Data are presented as mean ±SEM n = 4. Two-tailed Student’s t-test. **P < 0.01, ***P < 0.001, ns, not significant.

**Fig. S2.**
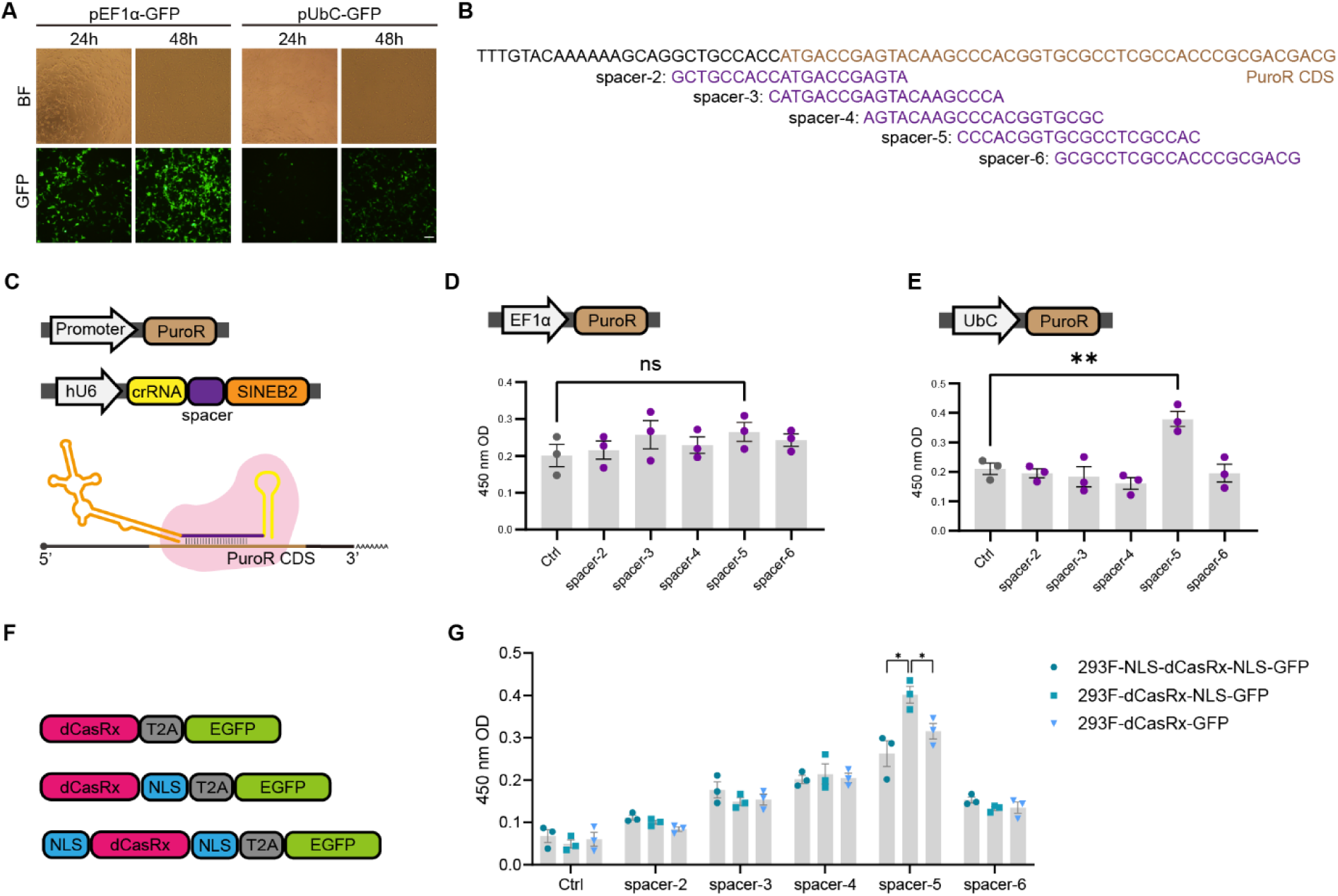
The design of other components of intracellular evolution, related to Fig 1. **(A),** The differences in EGFP expression between EF1α and UbC promoters at 24h and 48h. Scale bar = 200 µm. **(B), (D), (E),** Comparison of the differences in cell survival between the EF1α promoter and the UbC promoter at the same concentration of puromycin. NC (pcDNA3.1). Data are presented as mean ±SEM n = 3. Two-tailed Student’s t-test. **P < 0.01, ns, not significant. **(C),** The function of SINEB2 was verified by expression of PuroR. crRNA-SINEB2 was located on the mRNA of PuroR via dCasRx. **(F),** Schematic diagrams of dCasRx proteins with different nuclear localization sequences. **(G),** The result of CCK8. dCasRx with different NLS locates SINEB2 at different spacer sites. After transfection with the plasmid, cells were cultured in a medium containing puromycin and CCK8 was performed on the sixth day. NC (pcDNA3.1). Data are presented as mean ±SEM n =3. Two-tailed Student’s t-test. *P < 0.05.

**Fig. S3.**
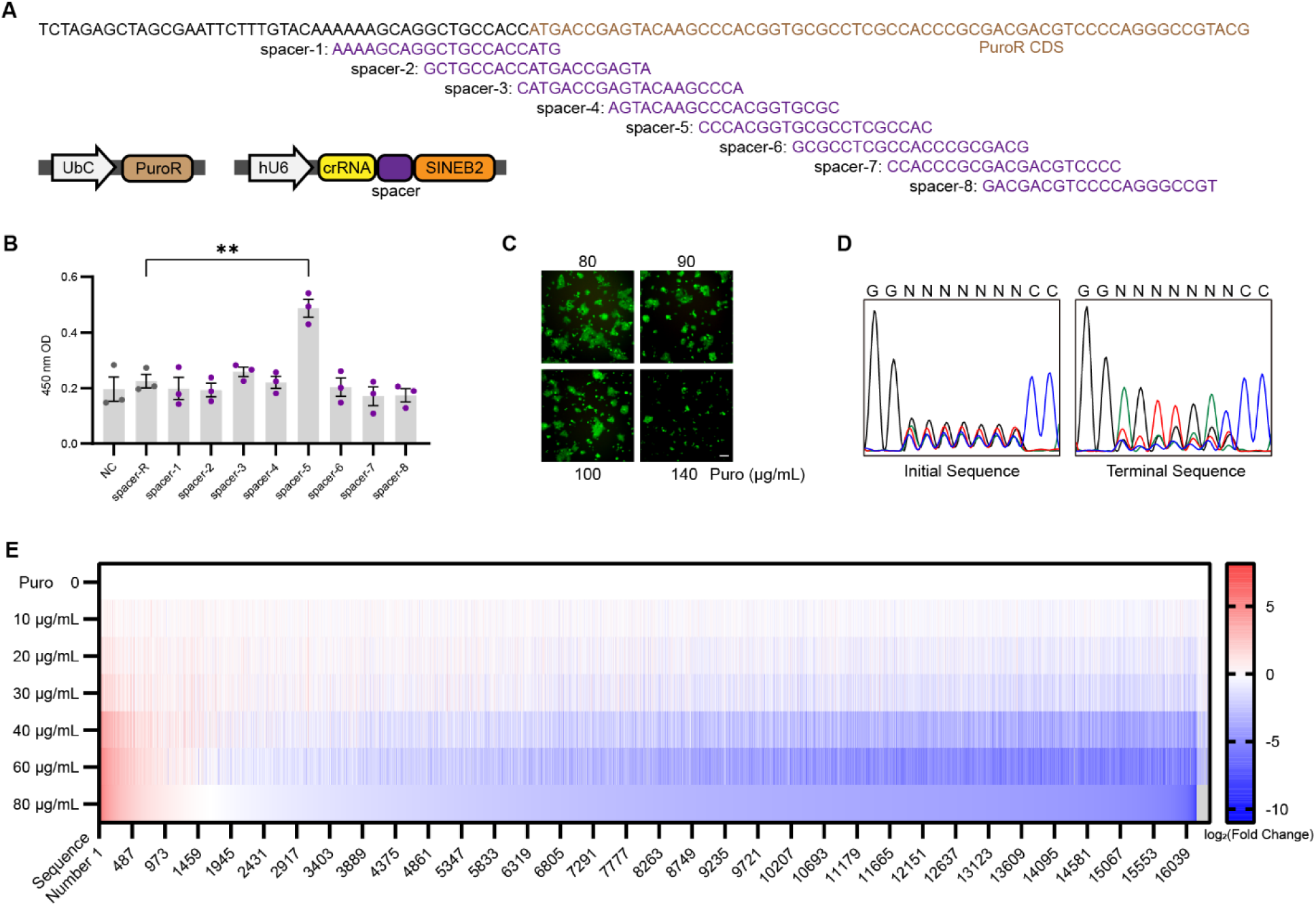
Characterization of intracellular evolutionary processes, related to Fig 1. **(A)**, Characterization of the selective effect of puromycin. The PuroR expression plasmid and the crRNA-SINEB2 transcription plasmid were co-transfected into 293F-dCasRx cells. Puromycin was added to the culture medium, and CCK8 was performed on the eighth day. **(B)**, CCK8 results of different Spacers. NC (pcDNA3.1). Data are presented as mean ±SEM n = 3. Two-tailed Student’s t-test. **P < 0.01. **(C)**, The proliferation of cells obtained at a concentration of 60 μg/mL puromycin in the medium containing higher puromycin. Scale bar = 200 µm. **(D)**, The results of sanger sequencing, including initial libraries and terminal libraries. **(E)**, The results of next-generation sequencing. The sequence is sorted in descending order by proportion of end point/ proportion of initial point. Compared with the proportion in the initial library, red represents the ascending sequence and blue represents the descending sequence.

**Fig. S4.**
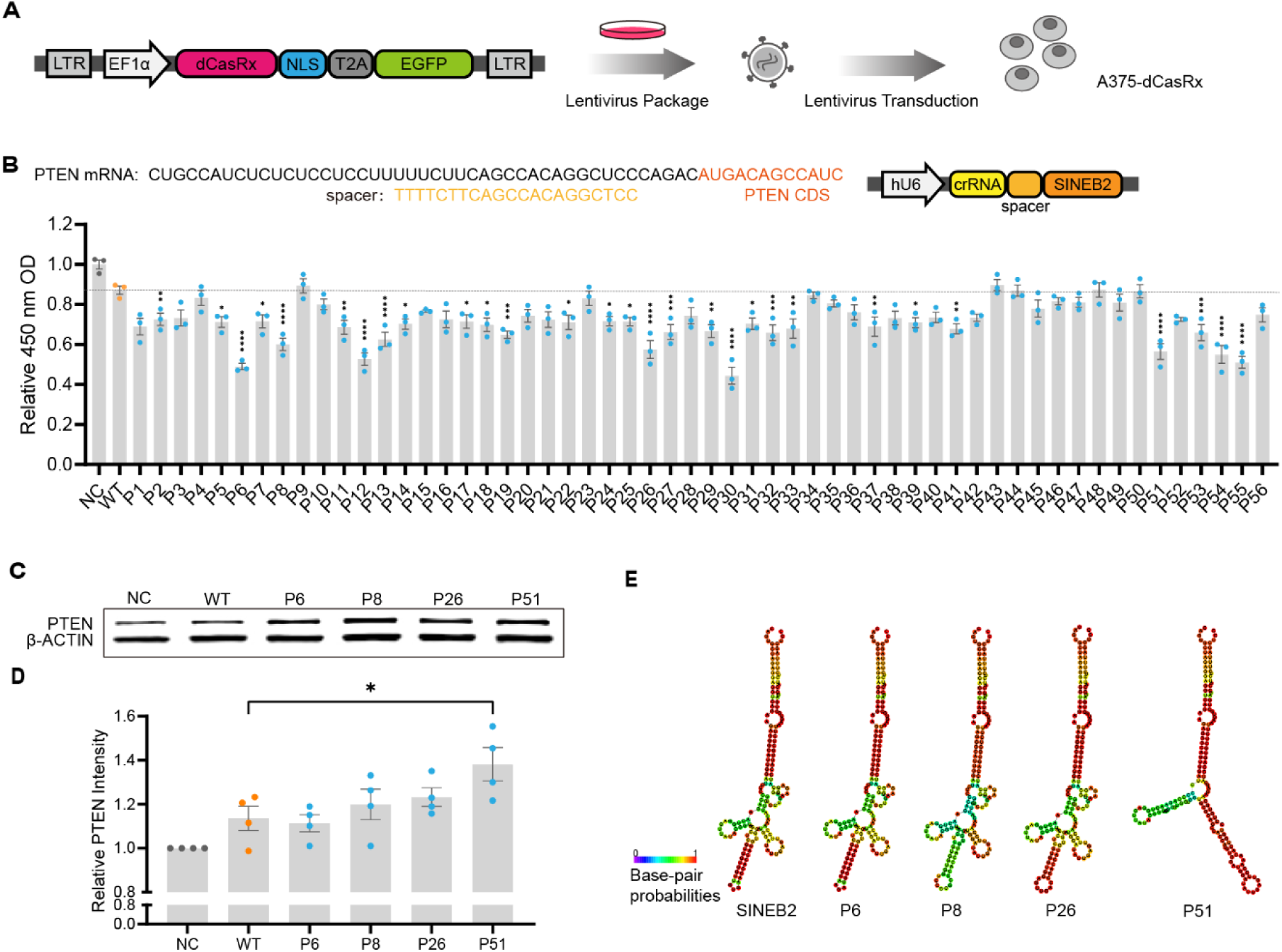
The sequence obtained through intracellular evolution, related to Fig 1. **(A),** Schematic diagram of the construction of the A375-dCasRx cell line. **(B),** Regulate the expression of endogenous gene. The function of the sequences obtained through screening was verified by the CCK8, in A375-dCasRx. The upper part indicates the plasmid structure and the localization of crRNA. PTEN is a tumor suppressor gene. Upregulation of PTEN can inhibit the proliferation of A375. NC (random sequence shuffled from SINEB2). Data are presented as mean ±SEM n =3. Two-tailed Student’s t-test. *P < 0.05, **P < 0.01, ***P < 0.001, ****P < 0.0001. **(C), (D),** Western blot data from the A375-dCasRx cells transfected with the plasmid of crRNA-sequences (SINEB2, P6, P8, P26, P51) at 48h post transfection. NC (random sequence shuffled from SINEB2). Data are presented as mean ±SEM n =4. Two-tailed Student’s t-test. *P < 0.05. **(E),** The secondary structure of RNA includes: SINEB2, P6, P8, P26, P51. The color represents the possibility of base pairing.

**Fig. S5.**
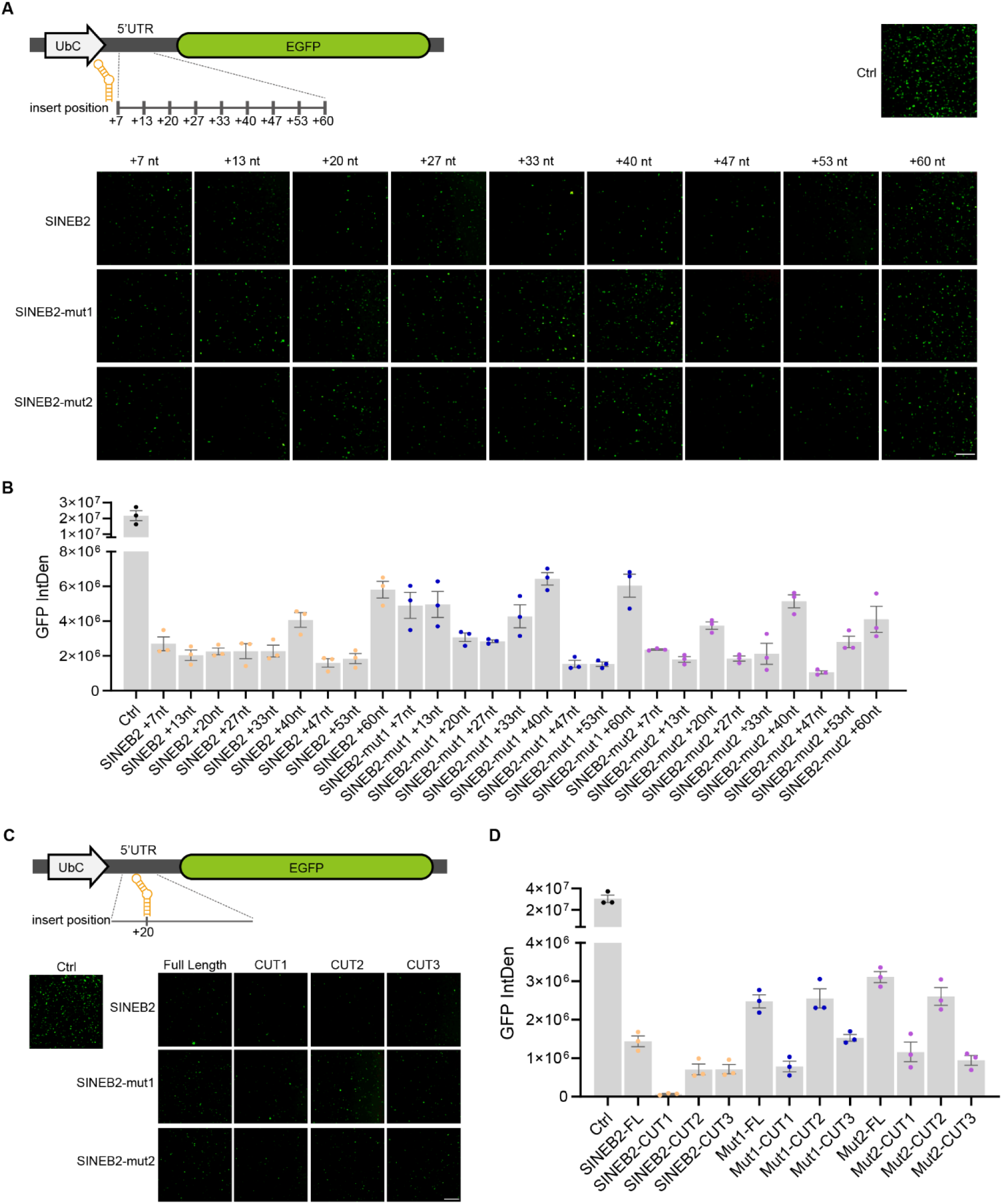
Engineering the 5’UTR with SINEB2 elements. **(A), (B),** The results of inserting SINEB2 elements or its two mutants at nine discrete sites within the 5’UTR of UbC-EGFP was evaluated by fluorescence spectroscopy in HEK293T cells transfected with the respective engineered plasmids. Scale bar = 1000 µm. **(C), (D),** The results of full-length SINEB2 element and its two mutants, as well as their three truncated derivatives, inserted within the 5’UTR of UbC-EGFP was evaluated by fluorescence spectroscopy in HEK293T cells transfected with the respective engineered plasmids. Scale bar = 1000 µm.

**Fig. S6.**
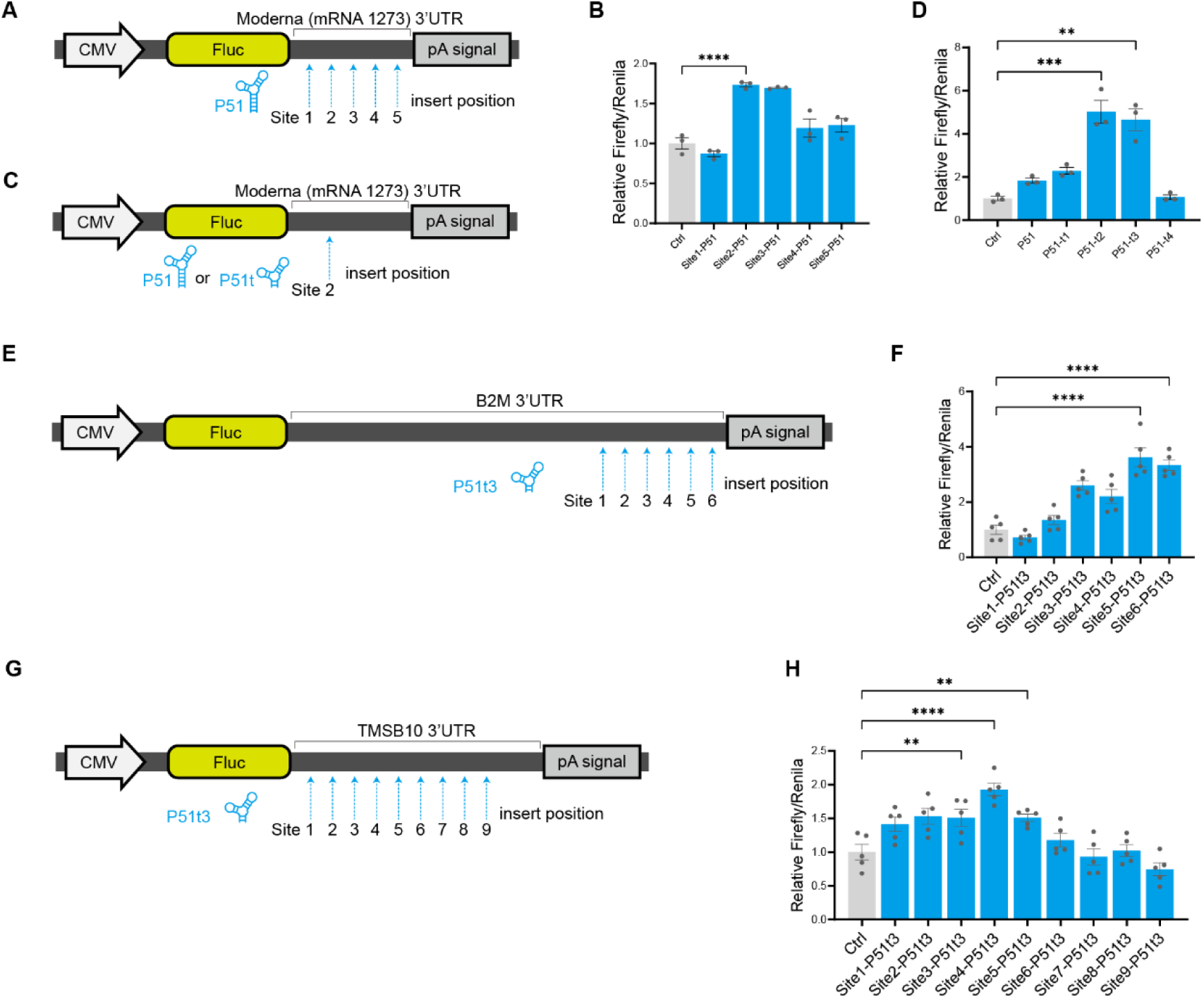
Wide applicability of P51t3 element in 3’UTR. **(A),** Schematic of P51 elements inserted at five sites in the CMV-Fluc reporter 3’UTR. **(B),** The results of P51 elements inserted at five sites in the CMV-Fluc reporter 3’UTR (Moderna mRNA-1273 3’UTR) was evaluated by dual-luciferase assay in HEK293T cells transfected with the respective engineered plasmids. Data are normalized to the CMV-Fluc reporter 3’ UTR lacking P51 insertions (Ctrl) and presented as mean ± SEM (n = 3). ****P < 0.0001, two-tailed Student’s t-test. **(C),** Schematic of P51 element and its four truncations inserted in the CMV-Fluc reporter 3’UTR. **(D),** The results of P51 element and its four truncations inserted in the CMV-Fluc reporter 3’UTR (Moderna mRNA-1273 3’UTR) was evaluated by dual-luciferase assay in HEK293T cells transfected with the respective engineered plasmids. Data are normalized to the CMV-Fluc reporter 3’UTR lacking P51 or P51-truncation insertions (Ctrl) and presented as mean ± SEM (n = 3). **P < 0.01,***P < 0.001, two-tailed Student’s t-test. **(E),** P51t3 is inserted at different sites of B2M 3’UTR. **(F),** P51t3 enhanced the translation of Fluc (B2M 3’UTR) in HEK293T. The results of the dual-luciferase reporter system from the HEK293T cells transfected with the plasmid at 48h post transfection. Data are normalized to the reporter 3’UTR lacking P51t3 insertions (Ctrl) and presented as mean ±SEM n = 5. Two-tailed Student’s t-test. **P < 0.01, ****P < 0.0001. **(G),** P51t3 is inserted at different sites of TMSB10 3’UTR. **(H),** P51t3 enhanced the translation of Fluc (TMSB10 3’UTR) in HEK293T. The results of the dual-luciferase reporter system from the HEK293T cells transfected with the plasmid at 48h post transfection. Data are normalized to the reporter 3’UTR lacking P51t3 insertions (Ctrl) and presented as mean ±SEM n = 5. Two-tailed Student’s t-test. *P < 0.05, **P < 0.01, ****P < 0.0001.

**Fig. S7.**
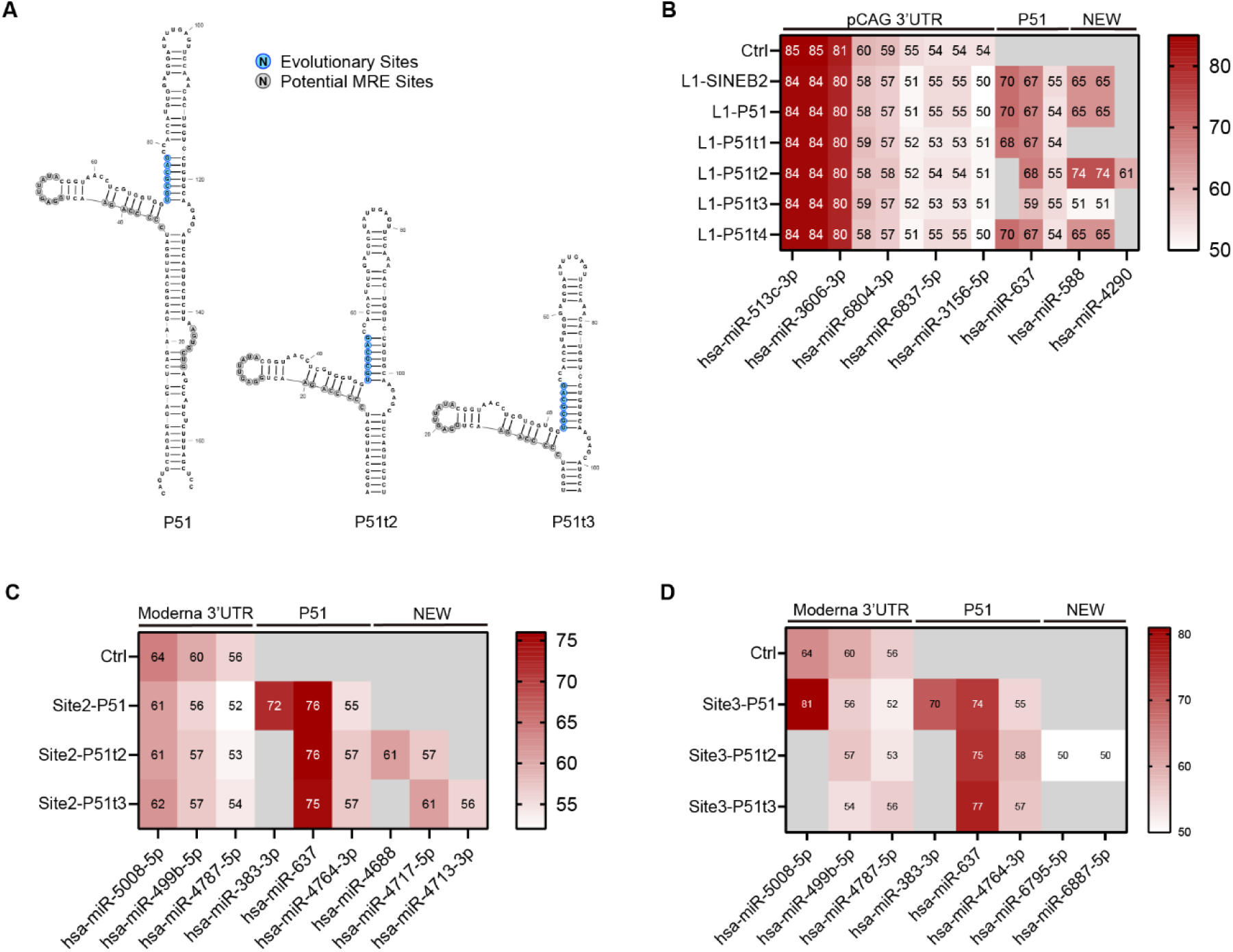
MRE analysis of the P51 element and its truncations inserted into the 3’UTR. **(A),** Structures of the P51 element and its truncations P51t2 and P51t3. Blue circles denote evolutionary sites; gray circles indicate potential MRE sites. RNA secondary structures were predicted using RNAfold WebServer and rendered using RNAcanvas. **(B),** Predicted MRE landscape of the Moderna mRNA-1273 3’UTR Site2 upon insertion of P51 or its truncations. Target scores are displayed as heat maps. MRE analysis was performed using miRDB. **(C),** Predicted MRE landscape of the pCAG 3’UTR L1 upon insertion of SINEB2 element or P51 element or P51-truncations. Target scores are displayed as heat maps. MRE analysis was performed using miRDB. **(D),** Predicted MRE landscape of the Moderna mRNA-1273 3’UTR Site3 upon insertion of P51 or its truncations. Target scores are displayed as heat maps. MRE analysis was performed using miRDB.

**Fig. S8.**
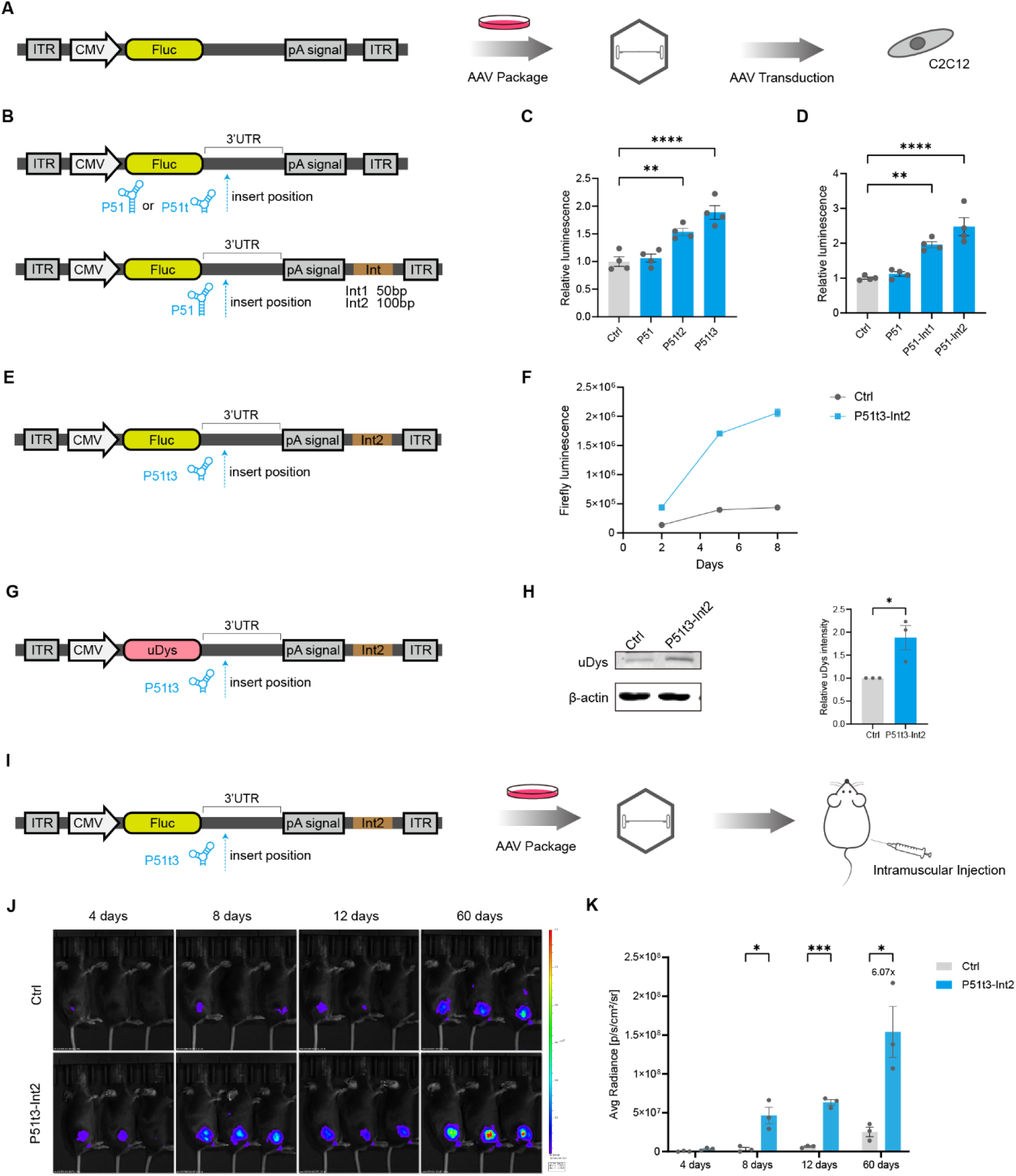
P51t3 element enhances the gene expression of the AAV vector. **(A),** Schematic diagr **|** m of AAV vector packaging and cell experiments. **(B),** Schematic diagram of the modification of the Fluc AAV 3’UTR. (i) Insert P51, P51t2 and P51t3 at the optimal position. (ii) Insert P51 at the optimal position. Insert additional random interval bases between polyA signal and downstream ITR. **(C),** P51t2 and P51t3 enhanced the translation of AAV vector Fluc in C2C12. Data are presented as mean ±SEM n =4. Two-tailed Student’s t-test. **P < 0.01, ****P < 0.0001. **(D),** Additional interval bases between polyA signal and downstream ITR enhanced the translation of AAV vector Fluc in C2C12. Data are presented as mean ±SEM n =4. Two-tailed Student’s t-test. **P < 0.01, ****P < 0.0001. **(E),** Schematic diagram of the modification of the Fluc AAV 3’UTR. Insert P51t3 at the optimal position. Insert additional random interval bases(100bp) between polyA signal and downstream ITR. **(F),** P51t3 and Int2 enhanced the translation of AAV vector Fluc in C2C12. Data are presented as mean ±SEM. **(G),** Schematic diagram of the modification of the uDys AAV 3’UTR. **(H),** Western blot data from the C2C12 cells transduced with the uDys AAV at 6 days. Data are presented as mean ±SEM n = 3. Two-tailed Student’s t-test. *P < 0.05. **(I),** Schematic diagram of AAV vector packaging and animal experiments. **(J), (K),** Exemplary in vivo luminescence images of mice treated with ctrl/P51t3-Int2 AAV at 4,8,12 and 60 days post AAV injection(10^11vg/site). Color scale of the heatmaps, Radiance. Luminescence was measured by integration of total radiance for each mouse. Data are normalized to the mice of Ctrl group. Data are presented as mean ±SEM n = 3. Two-tailed Student’s t-test. *P < 0.05.

**Fig. S9.**
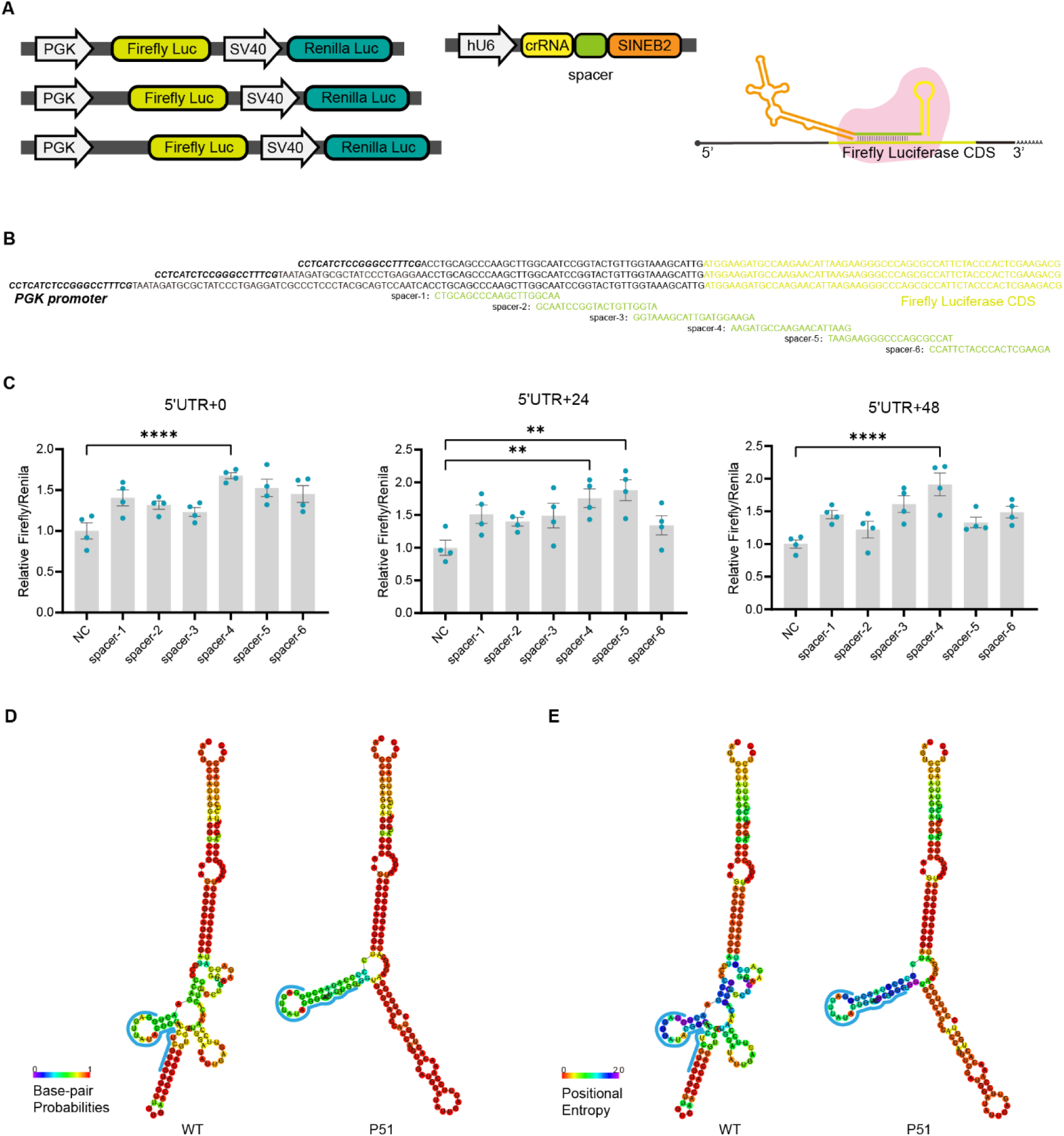
Analysis of the mechanism of P51t3 element, related to Fig. 7. **(A),** Schematic diagram of plasmid transfection. The 5’UTR lengths of Fluc plasmids were different. **(B),** Schematic diagrams of 5’UTR of different lengths and different spacer sites. **(C),** The results of dual-luciferase experiments with 5’UTR of different lengths. NC (pcDNA3.1). Data are presented as mean ±SEM n = 4. Two-tailed Student’s t-test. **P < 0.01, ****P < 0.0001. **(D),** Base-pair probabilities of WT and P51. The bases represented by the blue line were predicted to bind to 28s rRNA. **(E),** Positional Entropy of WT and P51. The bases represented by the blue line were predicted to bind to 28s rRNA.

**Table S1: Sequence of each element**

**Table S2: Information of plasmids, mrnas, primers and UTRs, etc**

**Table S3: Reagent and resource**

